# Critical Scaling of Novelty in the Cortex

**DOI:** 10.1101/2024.12.23.630084

**Authors:** Tiago L. Ribeiro, Ali Vakili, Bridgette Gifford, Raiyyan Siddiqui, Vincent Sinfuego, Sinisa Pajevic, Dietmar Plenz

**Author notes:** Correspondence: Dietmar Plenz, Ph.D., Section on Critical Brain Dynamics, National Institute of Mental Health, Porter Neuroscience Research Center, Rm 3A-1000, 35 Convent Drive, Bethesda, MD 20892.

## Abstract

The ability to detect and transmit novel events is essential for adaptive behavior in uncertain environments. Here, we investigate how holographically triggered, unanticipated action potentials propagate through the primary visual cortex of resting mice, focusing on pyramidal neuron communication. We find that these novel spikes — uncorrelated with ongoing activity — exert a disproportionately large influence on neighboring neurons, whose response scales as a power law (exponent ∼0.2–0.3). Even a few such spikes can recruit a large fraction of the local network, enabling robust decoding of perturbation origin despite high trial-by-trial variability and ongoing activity dominated by large activity fluctuations in the form of scale-invariant, parabolic neuronal avalanches. Simulations confirm this scaling to small, local perturbations aligns with the high susceptibility of complex systems near criticality. These results suggest that critical dynamics facilitate efficient transmission of novel signals, revealing a fundamental mechanism for cortical novelty detection.

## Introduction

Thriving in uncertain environments depends on the brain’s ability to detect and respond to unexpected events. In mammals, such events must be rapidly and effectively communicated across brain regions to support accurate evaluation and adaptive behavior^1^. In the cortex, the fundamental unit of this communication is the action potential generated by a single pyramidal neuron. Therefore, understanding how the brain processes unexpected events requires examining how novel action potentials — those that deviate from near-future predictions based on ongoing activity — are transmitted through the surrounding cortical network.

Neuronal spiking has been found to shape cortical activity and behavior. A spike in a pyramidal neuron from superficial layers (L2/3) of cortex of mice has been estimated to influence fewer than 1% of nearby neurons, to evoke strong local inhibition^2–5^, and in reptilian cortex to reliably recruit activity sequences^6^. Coordinated activity in small groups of L2/3 pyramidal neurons elicits spatially extended variable responses^7^ and can drive trained behavior via pattern completion^8,9^. Similarly, spike bursts in deep layers (L5/6) can elicit sensitized behavioral responses^10,11^ and alter global brain states^12,13^. These holographic perturbation studies in mice are in line with previous findings from cortical micro stimulation in primates highlighting the capacity of few spikes to significantly alter brain activity under specific conditions (for review^14^). However, a comprehensive framework to quantify the impact of action potentials on the spatiotemporal dynamics of cortical networks remains elusive.

In principle, novel spikes representing unexpected events should induce propagation across the cortex both broadly (to reach multiple functional domains for evaluation given their uncertain nature) and rapidly (to be able to act quickly). However, this dissemination of information is constrained by several factors. First, even in the resting state, the cortex exhibits substantial activity, with each neuron receiving thousands of action potentials every second.

Accordingly, individual neurons operate in a fluctuation-dominated^15^, high-dimensional^16^ regime that affects their ability to respond to and encode information from novel spikes^17^ or sensory inputs^18^. Second, the sparse synaptic connectivity between L2/3 pyramidal neurons —estimated at only 5–8% locally and diminishing sharply within 200 µm^19–23^ — further restricts the transmission of novel spikes^24^. Additionally, the weak and transient synaptic currents of these failure-prone connections in vivo^2,22^, coupled with strong local inhibition^25,26^, further reduce the potential influence of novel spikes introduced randomly into a pyramidal neuron in the awake state. Given these limitations, understanding how novel spikes are encoded requires a focus on the network’s broader dynamics rather than the activity of individual neurons or microcircuits within a particular experimental constraint. However, adopting this broader perspective presents a significant challenge due to the inherent complexity of cortical activity.

Research suggests that brain networks operating near a critical state exhibit heightened sensitivity to small perturbations^27,28^, which might enable the cortex to detect and process new information despite its complex internal activity and sparse topology. In this state, the system undergoes internal fluctuations while remaining responsive to small, local events that can propagate broadly, facilitating the recognition and dissemination of novel information. In the superficial cortical layers, these fluctuations manifest as “neuronal avalanches” — scale-invariant patterns of synchronized neuron activity characteristic of the awake state^29–31^. These avalanches align with theoretical predictions of critical dynamics. However, direct experimental evidence demonstrating that novel, localized information propagates within the neuronal avalanche regime and scales in line with predictions from critical dynamics is lacking, leaving a gap in understanding whether and how criticality supports the spread of novel information in the brain.

Using an all-optical approach, we holographically triggered spikes in individual L2/3 pyramidal neurons while monitoring activity of hundreds of nearby neurons in the primary visual cortex of awake mice. Our results demonstrate robust scaling of network responses during spontaneous, ongoing avalanche activity, with novel spikes rapidly generating widespread, yet transient information-rich activity across large cortical distances. Simulations show that this pattern of response emerges naturally in networks operating near a critical state, where small inputs can lead to large effects. Together, our findings suggest that critical dynamics play a key role in enabling the brain to efficiently process unexpected information.

## Results

### Holographically triggered spikes in a pyramidal neuron induce rapid, widespread excitation in superficial cortical layers

In awake, quietly resting adult mice, we holographically targeted single L2/3 pyramidal neurons (Target Cells, TC) in primary visual cortex (V1) that chronically co-expressed the calcium indicator jGCaMP7s and the opsin ChrimsonR (∼5 – 10 mW, 100 ms; ∼100 – 150 trials per TC) while simultaneously monitoring spiking activity from ∼100 – 300 nearby neurons using 2-photon imaging (Fig. 1a; 2PI, field of view: 450 µm × 450 µm; ∼50 – 140 µm cortical depth; temporal resolution: 22 ms; n = 6 mice; 16 experiments). Sparse co-expression, stimulation beam profiling, motion correction analysis, and anatomical response analysis confirmed select single neuron stimulation (Supplementary Figs. 1–5).

**Fig. 1.**
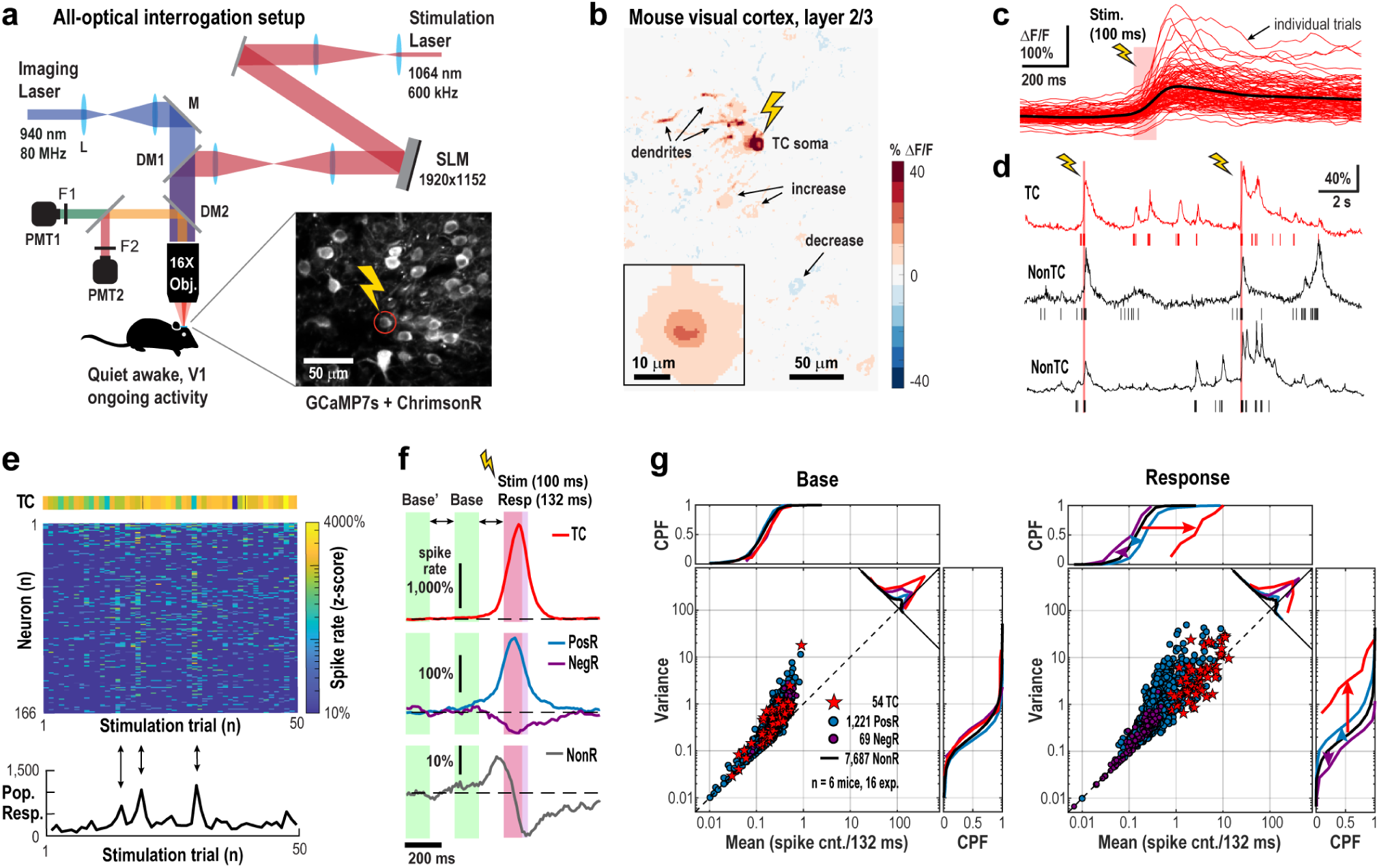
Single-neuron holographic stimulation drives widespread excitation in primary visual cortex of awake, resting mice. **a** Schematic of all-optical setup combining holographic stimulation and 2-photon imaging (2PI) in awake, resting mice (V1). (DM: dichroic mirror, M: Mirror, F: filter, PMT: photomultiplier, L: lens, SLM: spatial light modulator). Inset: Mean 2P image in L2/3 showing co-expression of jGCaMP7s and ChrimsonR; one pyramidal neuron (*circle*) was targeted (*flash*). **b** Stimulation (100 ms) of the target cell (TC) evokes relative fluorescence changes (ΔF/F) in soma, dendrites, and nearby neurons (*arrows*). Averaged over n = 150 trials. *Inset*: Mean across 54 TCs. **c** Variable ΔF/F responses to 100 stimulations (*red*) in a single TC with average in black. **d** Example ΔF/F traces (*lines*) and deconvolved spikes (*bars*) from the TC (r*ed*) and 2 responders (nonTC; *black*). **e** Z-scored spike counts in TC and surrounding neurons (n = 166) across 50 trials. *Bottom*: summed trial-wise z-scores showing variable network recruitment (*arrows*). **f** Average z-scored responses in TCs (n = 54), Positive Responders (PosR, n = 1,221), Negative Responders (NegR, n = 69) and Non-significant Responders (NonR, n = 7,687) from 16 experiments (6 mice). *Purple/Red bar*: response/stimulation window. Reconstructed protracted responses are due to deep interpolation (see also Supplementary Figs. 7 & 10). *Base’, Base* and *Resp* indicate respective time windows of analyses. **g** The response to single TC stimulation is largely excitatory, while maintaining variability above Poisson-level expectation (*dashed line*). Summary scatterplot of variance vs. mean spike count/132 ms for baseline and response (n = 6 mice, n = 16 experiments). Approximately 15% of neurons exhibit a significant increase in spike count from baseline (*PosR*), while fewer than 1% show decreased firing (*NegR*). For *NonR*, only CPF are shown. *CPF*: cumulative probability function.

Holographic stimulation of a single pyramidal neuron caused wide-spread, transient effects across the surrounding network (Fig. 1b – f; see also Supplementary Fig. 4). Within 132 ms (see Materials and Methods), approximately 15% of non-targeted neurons increased spiking (Positive Responders, PosR), while fewer than 1% showed decreased spiking (Negative Responders, NegR), despite high trial-by-trial variability (α = 2.5% significance threshold, one-tailed Welch’s *t*-test; see Materials and Methods). Ongoing, baseline activity was consistent across all groups (TC, PosR, NegR, and non-responders, NonR), with neurons exhibiting spike count variability well above Poisson expectations (Fig. 1g, base; n = 54 TC; n = 2,182 unique neurons, classifying into n = 1,221 PosR + 69 NegR + 7,687 NonR = 8,977 neurons due to multi-target experiments). Upon stimulation, PosR firing rates and spike count variance increased markedly remaining well above expectations for a Poisson process as reported previously^32^ (Fig. 1g; response).

### Novel spikes scale network responses without change in overall network variability

Holographically triggered spikes in TC neurons were classified as “novel” because they were not predicted by ongoing activity, as demonstrated by autocorrelation analysis. During the baseline period, ongoing spiking exhibited significant temporal order, with high autocorrelation across all groups (r ≈ 0.2, Fig. 2a, b; base’ *vs* base). However, during the response, the autocorrelation for TC neurons (base *vs* resp) dropped to nearly zero (r ≈ 0; p < 10^-3^, Wilcoxon rank sum), indicating that stimulation introduced spikes independently from the preceding baseline, disrupting the temporal organization of ongoing TC spiking. In contrast, temporal correlations remained stable for PosR and NonR, while the observed decrease in NegR autocorrelation reflected floor effects on suppressed low baseline activity (Fig. 2b).

**Fig. 2.**
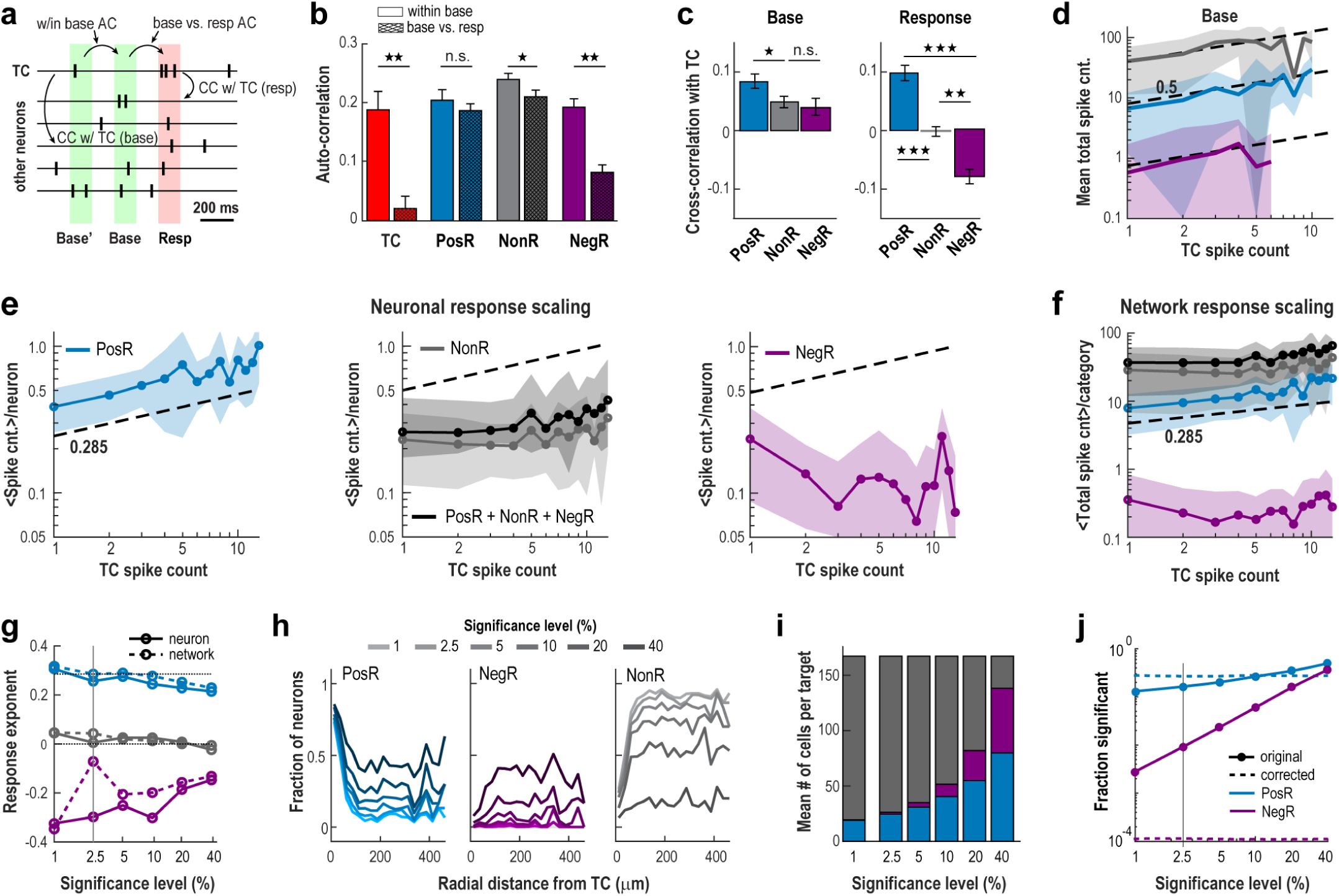
Scaling of cortical network responses to novel, holographically triggered spikes in individual pyramidal neurons in awake mice. **a** Schematic view of activity windows to calculate auto- and cross-correlations. **b** Holographically triggered TC spikes are novel, as evidenced by their zero autocorrelation with ongoing activity. In contrast, autocorrelations remain high for other neuron categories, except NegR, due to spike failure during the response. **c** Ongoing TC spike fluctuations positively correlate with all neuron groups. However, evoked TC spikes selectively correlate positively with 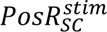, show no correlation with 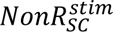 and correlate negatively with 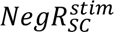. **d** Scaling for ongoing fluctuations in TC with respect to all other groups. Note limited range in spontaneous TC count fluctuations of <10 spikes/132 ms. Same color code as in panel e. **e** Robust scaling for novel spikes in 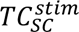 with respect to 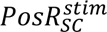 but not 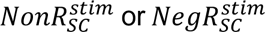. **f** Scaling of population spike count for each neuron category separately and combined (*black*). **g** Strong positive scaling for 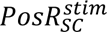 remains even when significance level α is greatly reduced (larger α). Change in scaling exponent as a function of α per neuron (*solid lines*) and population (*dashed lines*) for each group. **h** PosR are predominantly located within a 100 µm radius of the TC but also distribute throughout the network as α increases. **i** Relative number of neurons per category changes with α. Note that PosR and NegR make up the majority of the observed network at the lowest significance level at which robust scaling is still observed. **j** Fraction of significant responders as function of α. *Solid/dashed lines*: original/FDR-corrected data (see Materials and Methods). Vertical line indicates the level of α used for most analysis. *Shaded areas* (d – f) and error bars (b, c): SD across neurons. When comparing correlations (b, c), 1, 2 & 3 stars indicate p < 0.05, 10^-3^ & 10^-6^, respectively, using Wilcoxon rank sum test.

Spike count changes in TC in relation to other neuron categories exhibited distinct differences for baseline and response which we quantified using cross-correlations and scaling functions. During baseline, TC spike counts 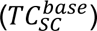 were significantly correlated with all groups (average cross-correlation <CC> ≈ 0.05; Fig. 2c) and followed a consistent scaling relationship:

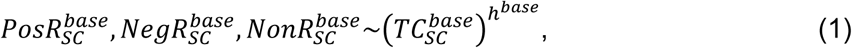

with *h^base^* ≅ 0.5 (linear regression; Fig. 2d). This scaling, which covered the majority of baseline fluctuations from 1 to ∼7 spikes/132 ms (transient firing rate increases of 8 – 53 Hz) was found both for the total spike count per neuron category and per-neuron spike count (Supplementary Fig. 6a).

During stimulation, only the excitatory response in PosR_SC_ showed scaling,

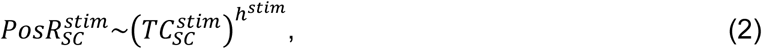

with a corresponding exponent of *h^stim^* ≅ 0.3 significantly lower compared to baseline (Fig. 2e, f; neuron: 0.26 ± 0.09; network: 0.28 ± 0.10; fit ± 95% confidence interval; see Materials and Methods; see also Supplementary Table 1). In contrast, NonR_SC_ exhibited no scaling, while NegR_SC_ showed negative scaling (see also Supplementary Fig. 6b). These holographic manipulations stayed within ongoing fluctuations in cortical firing as our 1–7 added spikes per 132 ms closely match the 1–7 spikes/132 ms seen during baseline (<20% of trials exceeded 7 spikes/132 ms). We also observed qualitatively similar scaling when measuring responses directly from the calcium transients (see Supplementary Fig. 6c).

The distinct scaling relationships between TC stimulation and the excitatory network response was mirrored in corresponding positive, zero, and negative cross-correlations between TC and the other neuron categories (Fig. 2c, right). Novel spikes preferentially scale with neighboring sub-networks composed of PosR that demonstrate a significantly higher baseline correlation compared to NonR and NegR (Fig. 2c, left). On average though, positive cross-correlation found for 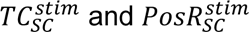 exhibited a relatively weak overlap in evoked and functional connectivity (see Supplementary Fig. 7; *R*^2^ = 0.018, linear regression; p < 10^-5^, for the *t*-statistic of the two-sided hypothesis test). Thus, scaling to novel spikes engages both established and alternative functional connectivities.

Scaling to novel spikes was robust to the significance level (α) employed. At ∼10 times higher α (less strict significance criteria), the scaling exponent *h^stim^* for 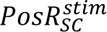 was still close to 0.2 (Fig. 2g). Despite our partial control of holographical spike generation due to variability from external (e.g., head movement) and internal (e.g., network inhibition) factors, these scaling relationships, nevertheless, derived a posteriori by averaging trials, confirm that driving a single pyramidal neuron distinctly affects neuronal populations compared to baseline. The scaling is also robust when running non-denoised (no Deep Interpolation) datasets (see Supplementary Fig. 8a).

The sparse synaptic connections between local L2/3 pyramidal neurons — estimated at only 5–8% and largely absent beyond 200 µm^19–23,33^ — suggest that among 200 neurons observed, less than 16 neurons are synaptically connected to a given TC. Indeed, PosR were consistently found near TCs within a ∼200 µm radius as reported previously^4^, in line with expectations for locally connected neighbors providing the strongest spike count increases. On the other hand, a substantial portion of PosR was distributed throughout the network (Fig. 2h) and this proportion increased as the significance threshold (α) was lowered (higher α). At the lowest significance level examined, nearly half of all recorded neurons showed significant positive responses, with an average scaling exponent of *h^stim^*∼ 0.22.

Recruitment of both PosR and NegR from the NonR pool (Fig. 2g–i) indicates that novel spikes are transmitted through the network via multi-synaptic pathways, triggering both increases and decreases in spiking, similar to what has been observed when stimulating groups of pyramidal neurons together^7^. When correcting for multiple comparisons^34,35^, we find that our significance threshold (α = 2.5) underestimates the number of significant PosR neurons by a factor of ∼1.7 times, while NegR become statistically non-significant. This demonstrates that novel spikes primarily drive a net excitatory response across the network (Fig. 2j). Notably, activating a single pyramidal neuron engages a large number of neurons spread across a wide area – an outcome that contrasts with what one would expect based on anatomical connections and synaptic physiology in activity regimes dominated by spontaneous fluctuations.

The robust scaling seen during TC stimulation may be partly due to a reduction in spiking variability, known to occur during sensory stimulation^36^ and often measured using the Fano factor (FF), which compares the variance to the mean of spike counts. We found that repeated holographic stimulation did lead to lower FF in TC neurons, consistent with more regular input. But for PosR and NonR neurons (defined at the original α = 2.5% threshold), FF did not change, even when we adjusted for differences in their average mean spike count^37^ (Fig. 3a, b). Likewise, there was no change in FF observed for PosR and NonR population responses (Fig. 3c). To test whether variability in TC spike triggering masked changes in FF for PosR and NonR, we analyzed subsets of trials with high or low TC spike count, for overall network activity and response timing. Yet, FF stayed consistently high and unchanged in both subsets under all conditions (Fig. 3d; Supplementary Fig. 9).

**Fig. 3.**
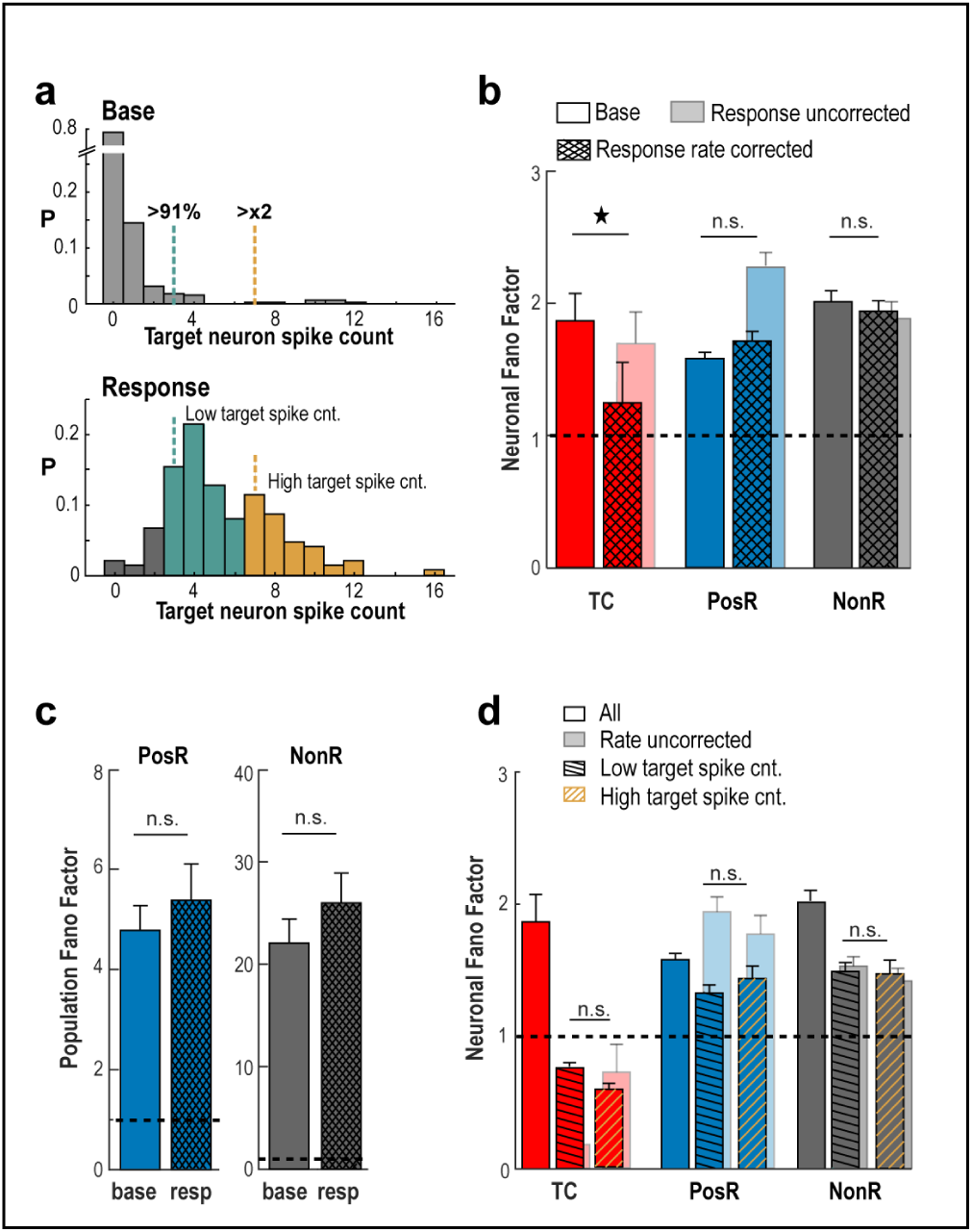
Network variability remains unchanged despite holographically triggered spikes. **a** Example TC with trials split by low vs. high spike count. *Top*: Base spike distribution defines 91st percentile and 2× threshold. *Bottom*: These thresholds are applied to classify stimulation trials. **b** The Fano Factor (FF) remains unchanged for PosR and NonR despite significant scaling in 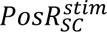 (α = 2.5%). *Transparent bars*: Mean spike count uncorrected FF (see Materials and Methods). **c** Population FF also shows no change across categories. **d** Summary statistics of mean TC trials separated by low and high target counts. Note the remaining high variance of PosR despite the large drop in TC variance. *Error bars* (b–d): SD across neurons. 1 star indicates p < 0.05, Wilcoxon rank sum test.

Our findings show that novel spikes from a single pyramidal neuron trigger strong, scalable responses in many nearby neurons, even though overall network activity remains dominated by fluctuations.

### Precise origin information from novel spikes spreads transiently across the cortical network

We next examined whether the broad network responses to novel spikes also encoded information about their point of origin. Across seven experiments, we stimulated up to 10 TCs in a semi-random sequence (each for 100 ms with 2–10 s between trials; Fig. 4a). We found that the origin of each novel spike event could be decoded with high accuracy from the activity of non-stimulated neurons (nonTC) using two machine learning approaches: XGBoost^38^ (XGB) and Random Forest^39,40^ (RF) (see Materials and Methods).

**Fig. 4.**
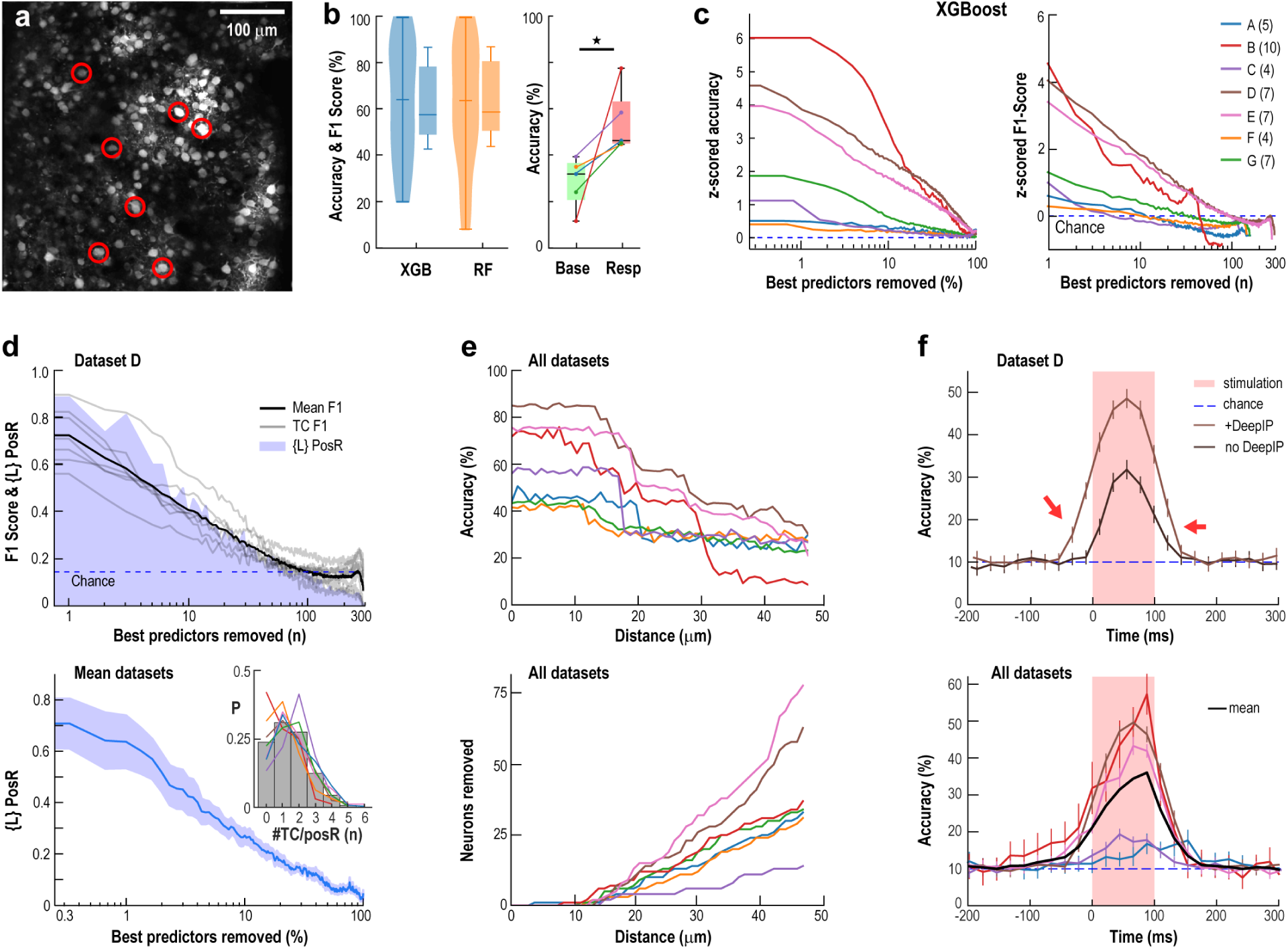
Origin of novel spikes propagates transiently and broadly in L2/3 of awake, resting mouse cortex. **a** Example field of view showing targeted TCs (circled) and surrounding network (mean baseline activity). **b** *Left*: Decoding TC identity using XGBoost (XGB) or Random Forest (RF) achieves highly above-chance accuracy and F1 scores (n = 7 experiments; chance: 10– 25%). *Right*: Accuracy when decoding from Base compared to Resp (see Materials and Methods; paired *t*-test, p = 0.012). **c** Distributed coding: a large fraction of the top-ranked neurons (10–70% or 10–100, by Shapley value) must be removed to reduce accuracy and F1 to chance (dashed lines; A–G experiments with number of TC indicated). **d** Likelihood (*L*) that dropped neurons are PosR. *Top*: *L* (*blue shade*) and F1 scores (*grey*: single TC; *black*: mean) for a single experiment. *Bottom*: average ± SD across experiments. *Inset*: Neurons can respond to multiple TCs— distribution of PosR counts per neuron. **e** Spatial restriction: excluding neurons by radius around TCs confirms localized stimulation. *Top*: Drop in overall accuracy when excluding all neurons within a certain distance from all TCs. *Bottom*: Number of neurons removed by this procedure. **f** Novel spike information is transient: Decoding accuracy rises above chance only when the response analysis window overlaps with stimulation (0–100 ms; *red shaded area*). *Top*: Single experiment ± Deep Interpolation (22-ms window). *Bottom*: All experiments (22-ms window, Deep-Interpolated).

Both algorithms achieved ∼60% decoding accuracy and F1-scores exceeded 90% for both algorithms when utilizing the full population of nonTC (Fig. 4b, left; Supplementary Fig. 10a). Decoding performance positively correlated with the number of trials in which the TCs were successfully activated (Supplementary Fig. 10b). For comparison, spontaneous spikes in TC occurring within 132 ms during ongoing activity (base) could also be decoded although with significantly lower accuracy (Fig. 4b, right; see Materials and Methods). Remarkably, it took the removal of 10–70% of the top-performing decoding neurons for accuracy to drop to chance levels (Fig. 4c). In a representative experiment (Fig. 4d), F1-scores of TCs dropped to chance levels after removing >70% of the best-encoding neurons. On average, 20–30% of the best-encoding neurons were sufficient to achieve high decoding accuracy, regardless of the algorithm used (Fig. 4c; Supplementary Fig. 10c).

Positively responsive neurons (PosR, defined at α = 2.5%) played a major role, contributing 30–60% of the decoding power (Fig. 4d). To confirm that decoding wasn’t driven by direct stimulation of nearby cells, we performed a spatial exclusion analysis. Excluding neurons within 10 μm of TCs had no effect on performance, but gradually removing more distant neighbors led to a steady drop in accuracy (Fig. 4e), confirming a spatially distributed response in the network to stimulation.

Individual neurons could significantly respond to stimulation from up to five different TCs, indicating overlapping functional subnetworks capable of encoding multiple spike origins (Fig. 4d, inset). Importantly, this information was short-lived — decoding accuracy dropped rapidly after the stimulation ended, in line with the brief activity profiles observed throughout the network (Fig. 4f, cf. Fig. 1f; analysis performed on a single frame – see Materials and Methods). We note that elevated decoding before the stimulus onset was attributed to temporal averaging (132 ms window) and Deep-Interpolation, which enhances signal quality but slightly blurs response timing (see Supplementary Fig. 10d). Qualitatively similar results were obtained using the calcium transient directly (Supplementary Fig. 10e).

Together, our results show that superficial layers of cortex can briefly and reliably encode when and where a novel spike originates. This supports the existence of a robust, distributed coding mechanism for processing unexpected inputs in real time across the network.

### Novel spike encoding persists across fluctuation-dominated, network state of parabolic avalanches

Our findings so far demonstrate the remarkable impact of just a few action potentials amid the thousands of spikes generated every second in the awake brain’s cortical neighborhood. Given our 0.45×0.45 mm field of view and ∼170 imaged neurons (*cf*. Fig. 2i), we estimate our sampling to be ∼5% of this neighborhood^19^]. On average, we observed ∼23 PosR per TC (α = 2.5%), which would translate to more than 450 neighboring neurons significantly *increasing* their firing within tens of milliseconds in response to our holographic stimulation. This estimate, however, is likely highly conservative as increasing α (lowering the significance criteria) further increases the number of neighboring neurons with detectable changes in spiking, as confirmed by our decoding results. This adjustment increasingly recruits both positive and negative responses from the pool of previously non-significant responders (it is important to note that we are likely heavily underestimating the number of NegR^25,26^, given the constraints to detect significantly lower spike counts from neurons that fire very sparsely; see Materials and Methods).

Furthermore, for many neurons, the spike count variability during novel spike encoding exceeded what would be expected from a simple Poisson process (Fig. 1), with Fano factors comparable to those observed during ongoing spontaneous activity (Fig. 3). This high variability extended across neuronal subtypes, including positively responsive neurons (PosR). To better characterize this fluctuation-dominated regime, we analyzed its statistical properties in the context of novel spike encoding.

Holographic stimulation frequently occurred during periods of heightened ongoing activity (Fig. 5a), defined by continuous population spiking above a low threshold. These active periods followed heavy-tailed distributions in both size (total spike count) and duration, consistent with previously described neuronal avalanches^29,41^ (Fig. 5b). After correcting for subsampling using temporal coarse graining, we found that the average avalanche size scaled quadratically with duration — a key feature of parabolic avalanches^29,41,42^. These parabolic events included avalanches lasting up to ∼0.7 seconds, which accounted for 87% of all avalanches and about 20% of spikes during spontaneous activity (Fig. 5c, d). Longer avalanches (>1 s) showed near-linear scaling, suggesting activity too widespread to be fully captured within the imaging field of view^29,41,43^.

**Fig. 5.**
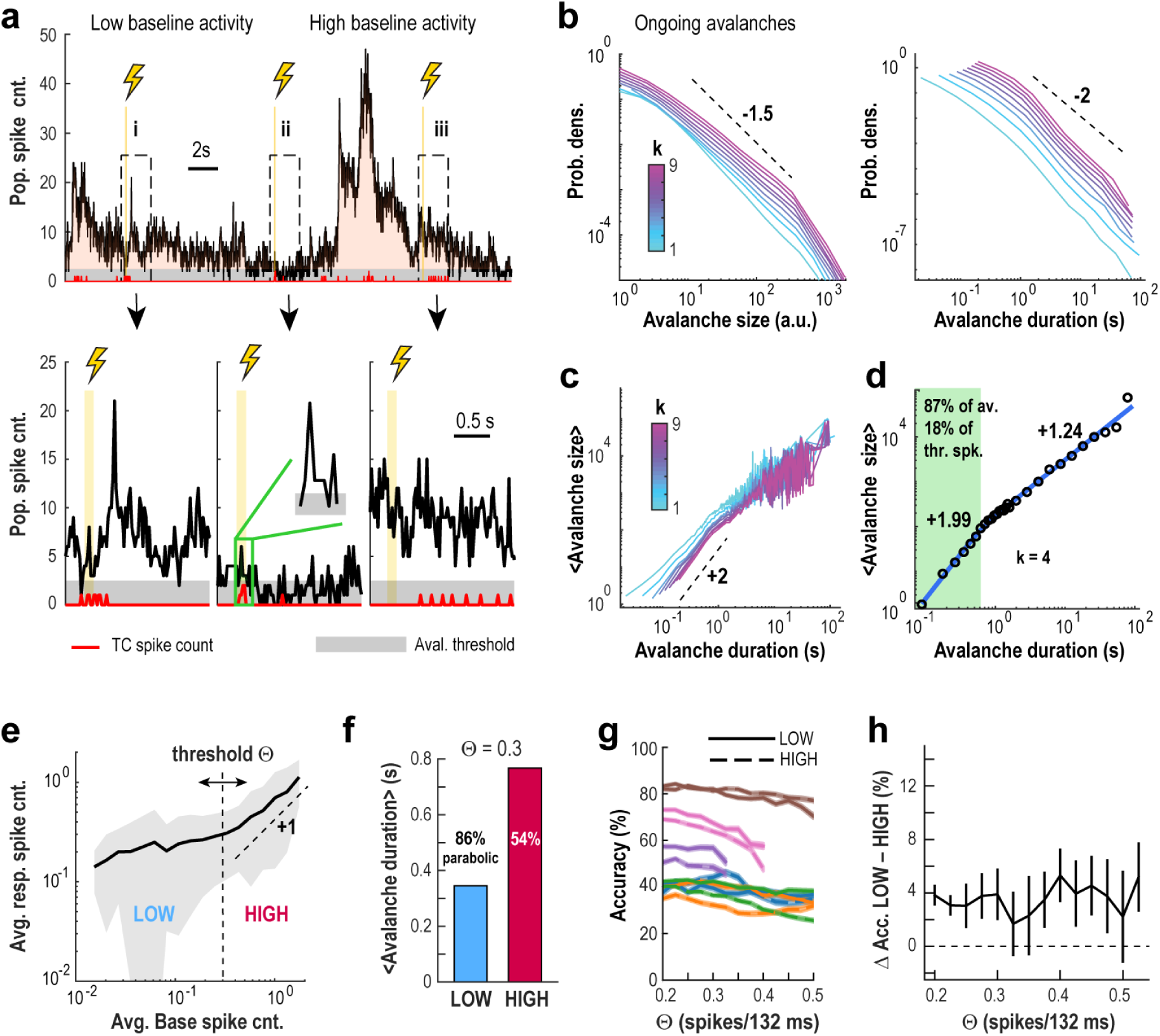
High decoding accuracy of novel spikes occur during scale-free avalanche dynamics despite highly fluctuating levels of baseline activity. **a** *Top*: Population activity in visual cortex of the awake mouse exhibits large fluctuations during quiet resting. *Bottom*: Zoomed in view of population activity in response to 3 (i – iii) holographic stimulations of a single TC. *Vertical lines*: TC stimulation. *Red line*: TC spike count. *Grey stripe*: Population threshold for avalanche detection. Shown is the population spike count (excluding TCs) per 22-ms frame. **b** Suprathreshold contiguous fluctuations in population activity exhibit the characteristic hallmarks of neuronal avalanches. Avalanche size (*left*) and duration (*right*) distributions as a function of the temporal coarse-graining factor *k*. **c** Scaling of avalanche dynamics as a function of temporal coarse-graining *k*. **d** Approximately 87% of avalanches are parabolic in nature, with durations <0.7 s, containing about 18% of thresholded spikes. Summary scaling of parabolic avalanches is shown for durations up to ∼0.7 s (*k* = 4). **e** Mean response spike count scaling as a function of mean baseline spike count for non-TC neurons. This analysis reveals a LOW baseline activity regime (mean baseline spike count < ∼0.3), in which responses are mostly independent from base activity, distinct from a HIGH baseline activity regime in which response activity can be predicted from base. **f** Mean avalanche duration is more than double for avalanches that start during LOW base trials compared to HIGH. 86% of all LOW avalanches are parabolic (scaling of χ = 2), whereas ∼54% of HIGH avalanches are parabolic. **g** Decoding accuracy remains largely unchanged when obtained using only LOW or HIGH baseline activity regimes, regardless of the threshold used to define the two regimes. Overall accuracy for LOW base trials (*solid lines*) and HIGH base trials (*dashed lines*) for each dataset (same color code as in Fig. 4), as function of the threshold used to define LOW and HIGH base trials. **h** Difference in accuracy for LOW and HIGH trials is small (with LOW trials having slightly higher accuracy), regardless of the threshold used to separate them.

Despite these dynamic and large-scale fluctuations, decoding the origin of novel spikes remained highly accurate across avalanche activity regimes. When baseline activity was low (θ < 0.3 spikes/132 ms), stimulus-evoked spike counts scaled modestly; when baseline activity was higher (θ > 0.3), the scaling approached a value of 1, indicating strong coupling between pre-stimulus and evoked activity (Fig. 5e). This allowed us to classify activity into low- and high-baseline regimes, both of which prominently featured parabolic avalanche dynamics (Fig. 5f).

Importantly, decoding accuracy differed by less than 4% on average between these regimes, regardless of the threshold used (Fig. 5g, h), and response scaling remained consistent across both states (Supplementary Fig. 11a).

These results suggest that cortical networks in vivo operate within a fluctuation-dominated regime where variable responses to local perturbations scale in line with expectations from critical dynamics^28,44^, which we further explore in the following section.

### Model simulations reveal observed experimental perturbation responses to be in line with those from critical networks

To study how networks with varying internal excitability respond to local stimulation, we simulated a two-dimensional network of 90,000 excitatory units with local, radial connectivity P_conn_(r) and three-state dynamics (rest, active, refractory) (Fig. 6a). Analysis focused on a central field of view of 456×456 µm^2^ with 1,440 neurons, including 10 target cells (TCs), to mimic experimental access and reduce edge effects.

**Fig. 6.**
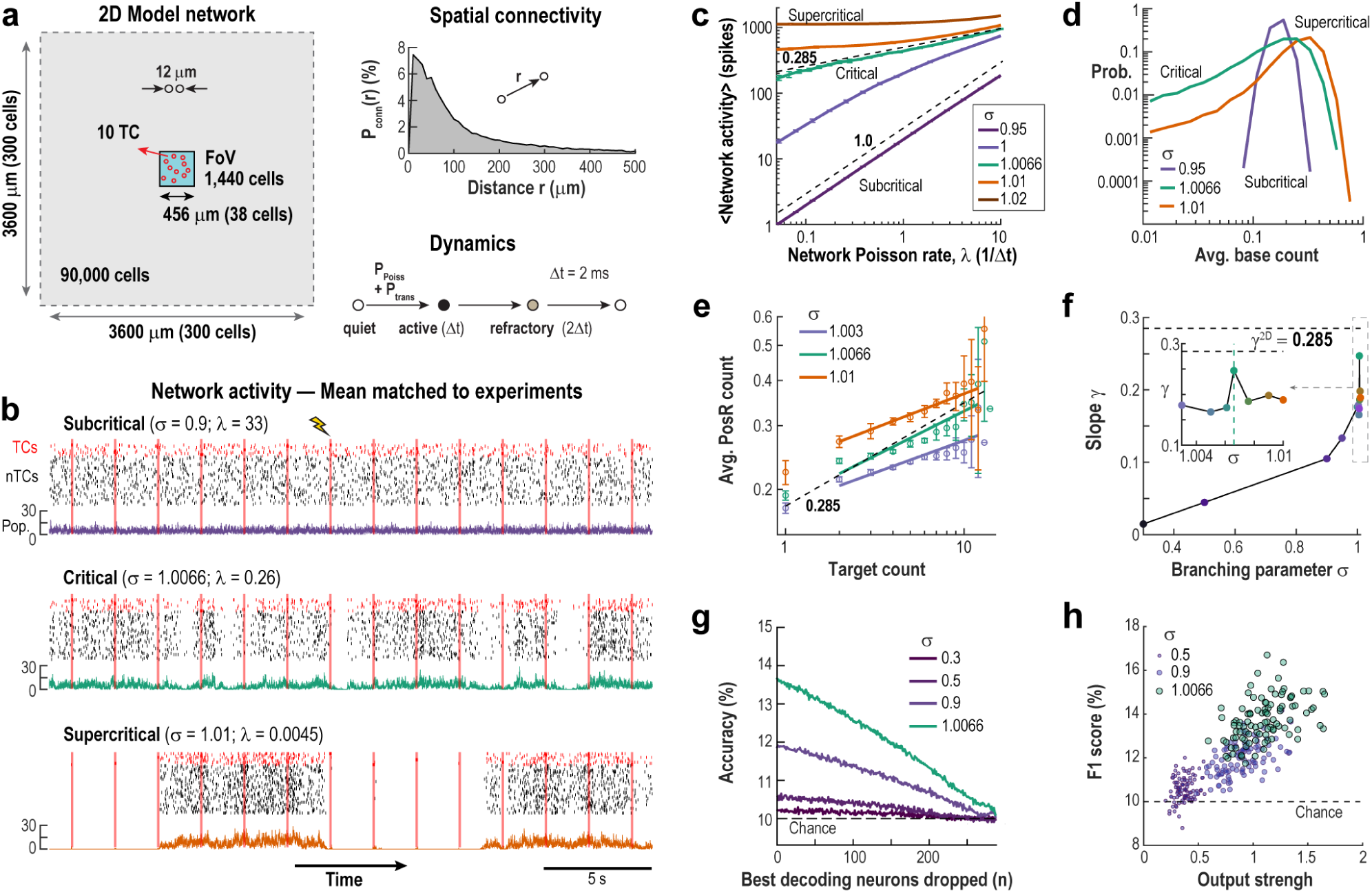
Simulated networks near criticality replicate experimental response scaling and decoding. **a** Simulation setup: A 2D network of 90,000 neurons with local spatial connectivity (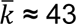 connections/neuron) receives external Poisson input. A central 1,440-neuron region (FoV; 456 x 456 µm^2^) with 10 TCs was analyzed to mimic experimental conditions and reduce edge effects. An external Poisson drive with fixed rate and active neighbors with fixed probability allow neurons to sequence through quiet, active, and refractory states. **b** Network activity under subcritical, critical, and slightly supercritical conditions shows TC (*red*) and non-TC (*black;* 50 neurons randomly selected) spiking during repeated TC stimulation (*red bars*). *Bottom*: Color coded summed activity of from the whole FoV. **c** Response scaling as a function of spike transmission probability (*P_trans_* = σ/⟨k⟩; *σ* is the branching parameter) reveals a power law (exponent ∼0.285) at the critical point (*magenta*). Subcritical (*blue*) and supercritical (*green*) regimes show linear and self-sustaining behavior, respectively. **d** Distribution of mean spike counts during baseline. Note decrease in activity fluctuation range during baseline for subcritical systems (*blue*). **e** Mean response of PosR as function of TC spike count. Close to criticality, response curves follow a power law profile similar to the one observed in experiments (cp. Fig. 2e). **f** Power law exponents for the response curves of PosR for varying branching parameter. Note that the exponents approach the value observed in experiments (∼0.28) close to criticality. **g** Decoding accuracy is highest close to criticality and drops to chance only after removing a very large fraction of the best predictors. Accuracy as function of number of best decoding neurons dropped for varying branching parameter. **h** F1 scores increases with TC’s (*circles*) output strength. Output strength is measured by the sum of all probabilities that a single spike in the TC, in a quiet network, would propagate to each of its direct neighbors, secondary neighbors, etc. up to order 5. It estimates the expected number of spikes that each spike in the TC would generate on average, ignoring refractory period and potential collisions (see Materials and Methods).

Each resting unit could be activated either by an external Poisson input at constant rate P_Poiss_, or by an active neighbor with transmission probability P_trans_. The system exhibits dynamics of the directed percolation universality class, with a phase transition centered around a critical state — between a subcritical phase (where activity fades without input) and a supercritical phase (where activity can self-sustain).

Near criticality, the branching parameter σ — the average number of near future downstream spikes per originating spike — is close to 1. We define 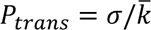, where 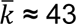 is the average number of connections per unit. Adjusting σ, moves the system in and out of the critical state. Empirically, the critical point was near σ = 1.0066, slightly above 1 to compensate for propagation loss (e.g. spike collision) in the network.

Figure 6b shows example traces of ongoing network activity under subcritical, critical, and slightly supercritical conditions, highlighting both the increase in overall activity and the emergence of larger fluctuations as the network approaches criticality. This shift is quantified in Fig. 6c, where the relationship between network activity and external drive (P_Poiss_) reveals the critical transition. In the subcritical regime (σ << 1), inputs spread weakly, and network activity increases linearly with external drive, yielding a slope γ ≍ 1 (see Refs.^45,46^). In the supercritical regime (σ > 1.0066), even low input leads to network saturation, resulting in γ ≍ 0. At the critical point, the system produces scale-invariant responses – external spikes can trigger cascades of all sizes – producing a power-law relationship with an intermediate slope 0 < γ < 1. For our 2D network, the predicted critical exponent is γ = 0.285 (see Ref^28^), as shown in Fig. 6c (magenta curve). To match experimental conditions, we adjusted P_Poiss_ in each simulation to reproduce average baseline activity observed in our in vivo experiments.

As shown in Fig. 6d, the distribution of ongoing firing rates is narrow in subcritical conditions but broadens near the critical point, reflecting a fluctuation-dominated regime. When measuring the response on PosR as function of TC spike count, we observed power-law scaling, with the slope approaching the one obtained experimentally (*h^stim^* ∼ 0.285) at criticality (Fig. 6e, f). Interestingly, when considering only trials in which the network had a low amount of activity preceding the stimulation, the slope of the scaling responses crosses the value 0.285 close to criticality, whereas considering only trials with a high amount of activity we observe that the highest slope happens at criticality, albeit at a smaller value, as in the experiments (Supplementary Fig. 11b, c). Importantly, classification accuracy improved sharply from subcritical to critical regimes and decayed gradually as increasing numbers of neurons were removed (Fig. 6g), again matching experimental observations. Furthermore, decoding success was higher for targets that were more well-connected to the network (output strength estimates the expected number of spikes each TC spike would generate in a quiet network; see Materials and Methods) and was maximized at criticality (Fig. 6h).

Despite higher activity fluctuations, this critical regime maximized the number of positive responders (PosR) to TC stimulation (Fig. 7a). As in the experimental results, the fraction of cells that are PosR is maximal around TCs and decay over larger distances (Fig. 7b). At criticality, about 93% of neurons receiving direct input from TCs were classified as PosR. However, only about 16% of all PosR had a direct connection from TCs, showing the importance of multi-synaptic pathways in the critical network. This reveals a unique “mixing” effect at criticality: while a TC’s average output strength is higher, its functional spread—the range of PosRs it engages—becomes much broader, reflecting the greater diversity seen in critical states (Fig. 7c).

**Fig. 7.**
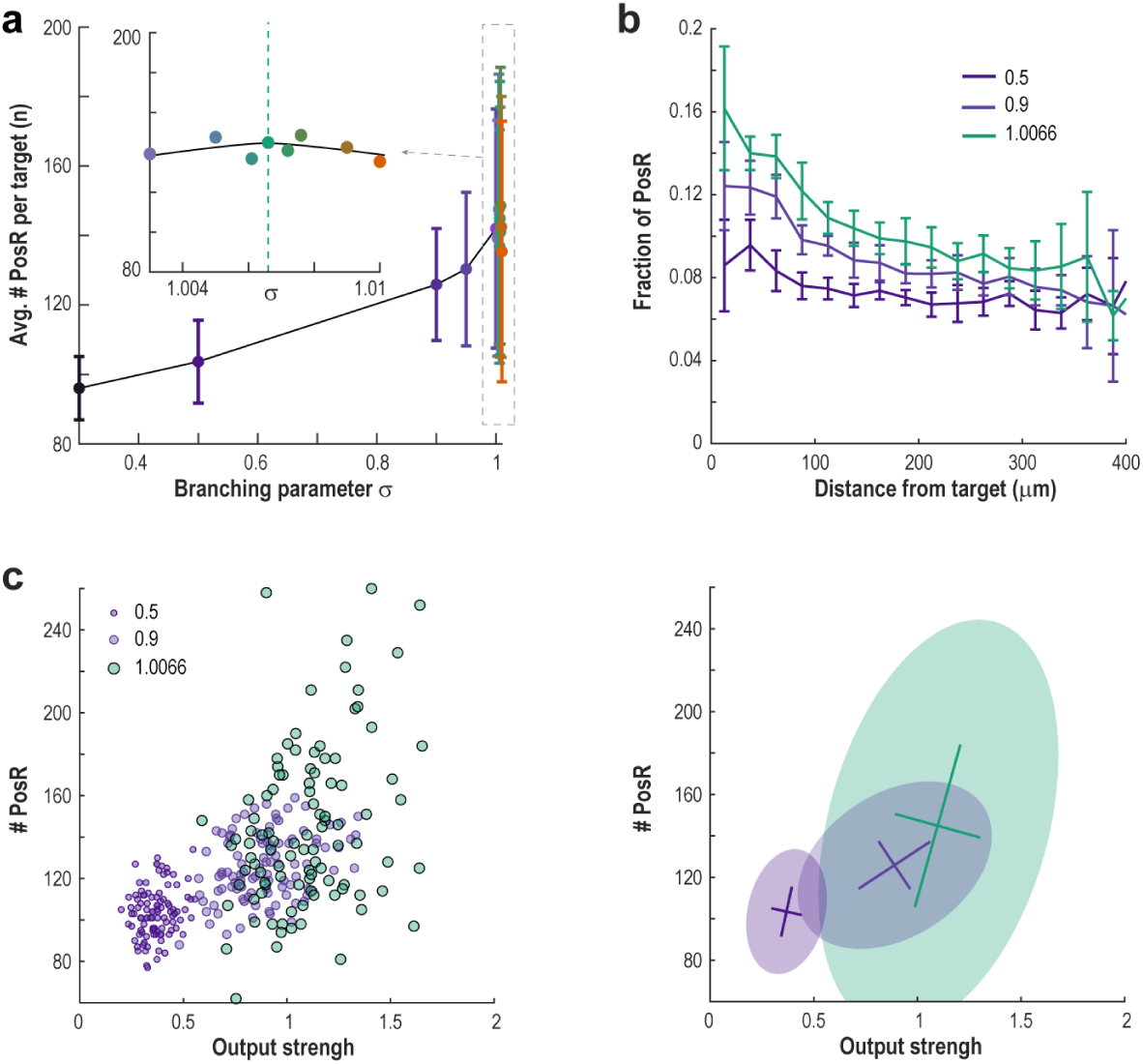
PosR are close and relate to output strength of TCs. **a** The mean number of PosR per target peaks close to the critical point of the network. **b** Average fraction of neurons that are classified as PosR as function of distance from TC. As expected, the fraction is highest close to TCs and decreases as the network deviates from criticality. **c** Number of PosR of each TC (*circles*) as a function of the TC’s output strength. *Right*: Covariance matrix ellipses (*shaded area*; 2.5 s.d.) and center of mass (*lines*; mean ± s.d.) for the scatter plot on the left.

To further examine how the network’s dynamical regime affects information spread, we measured the number of PosR for each TC in the subcritical network (σ = 0.5) and compared it to that of the critical network (σ = 1.0066). Even though the two networks have the same structure, the number of PosR for each TC is uncorrelated between them (Supplementary Fig. 12). This means that measuring a TC’s effect under subcritical conditions (lower excitability) gives little or no indication of its impact near criticality.

Together, these results strongly support the idea that novel inputs are most effectively propagated in networks operating near criticality, where activity is governed by fluctuations and scale-invariant dynamics.

## Discussion

Efficient communication of novel neural signals is vital for processing surprises and guiding future actions. Using an all-optical approach allowed us to precisely induce action potentials holographically in identified neurons and simultaneously measure its effect in the network at cellular resolution and high fractional sampling of the neuronal population in the awake, quietly resting mouse. Alternative perturbation approaches, such as extracellular microstimulation lack spatial selectivity^47^, or as in the case of intracellular patching^2^, are limited in capturing the network response. Here, we demonstrated in awake animals that introducing spikes in a single L2/3 pyramidal neuron rapidly activates widespread, net excitatory responses in the surrounding cortical network, encoding the novel stimulus’ origin. Remarkably, these responses scaled robustly and were efficiently decoded amidst ongoing, large fluctuations in network activity organized as scale-invariant, parabolic neuronal avalanches. Our scaling exponent of ∼0.2–0.3 in response to novel spikes was captured in a 2-dimensional network with critical dynamics allowing for highest decoding accuracy and engaging large neighborhoods in encoding stimulus origin. This high susceptibility and scaling to local perturbations^27,28^, suggests critical dynamics as a cornerstone of cortical processing, offering key insights into brain function.

Our findings challenge a longstanding discussion on cortical coding fueled by the intricate statistical anatomy of the cortex and its synaptic connections. Pyramidal neurons in associative layers establish few synaptic connections with their neighbors^19–23^. These excitatory connections are relatively weak, prone to failure, and decay within milliseconds^22,48^. The combination of sparse and weak connectivity makes it unlikely that a spike in a single neuron reliably triggers activity propagation in the surround network. To address this, most cortical codes rely on activity patterns that amplify synaptic impact on postsynaptic pyramidal neurons.

For instance, coincident activation of numerous synapses from diverse neurons converging onto a postsynaptic target enhances the likelihood of activity propagation, favoring coding models based on local synchrony, such as synfire chains^49,50^, waves^51,52^ or phase-locked oscillations^53,54^. Alternatively, repeated or burst firing of individual neurons^12,55^, strong synaptic connections^56^, subthreshold priming by background activity^15,57^ supported by subcellular amplification mechanisms such as short-term synaptic facilitation^58,59^ and non-linear dendritic processing^60^ can dynamically increase individual connection strength supporting rate-based coding.

Our findings do not support a dichotomic separation into synchrony vs. firing rate separation to secure propagation of neuronal activity (see also^24,61^). We show robust scaling of novel spikes within the cortical neighborhood from single to multiple action potentials delivered within 100 ms without an obvious threshold above which propagation is facilitated. This suggests that neither tight synchrony nor repeat firing is strictly necessary for effective encoding of novel spikes^62^. This conclusion is in line with findings that spike counts over 100 ms predict behavior independent of considerations of synchrony at the millisecond level^63^.

By emphasizing the sufficiency of a few action potentials in cortical codes, our results thus challenge existing paradigms and highlight a minimalist yet robust mechanism for cortical information processing supporting the prominence of sparse, single action potential firing found in vivo for associative layers^64^. Furthermore, the recruitment of significant responders is most likely composed of a mixture of direct (excitation, inhibition) as well as indirect (disfacilitation, disinhibition) local circuit interactions^4,7^, while preserving the general scaling relationship (cf. Fig. 2). Simulations have found that in such a fluctuation-dominated but relatively balanced regime, a reduction in membrane conductance due to increased synaptic bombardment still enables neurons to generate few spikes in response to rapid changes in synaptic input^17^.

The importance of critical dynamics for sensory cortical networks has been argued from the point of view of optimal decoding of stimulus intensities through maximization of the dynamic range^46^. In that context, a critical network is expected to produce response-input curves that scale as a power law, with exponents that are typically much smaller than one^28^. These low exponents allow the network to present distinct responses for a wide range of stimuli. On the other hand, the difference between these distinct responses gets smaller as exponents approach zero, indicating the possibility of an optimal value depending on the necessity of having a larger dynamic range versus the need for a more precise differentiation between responses. The study of psychophysics has demonstrated that psychological value scales with stimulus intensity as a power law with <1 exponents, a result known as Stevens’ law of psychophysics^65^. Therefore, it has been argued that the origins of Steven’s low exponents for response curves might originate precisely at this critical networks’ ability to amplify low-intensity stimuli and thus respond to a wide range of stimulus intensities^66^. We were able to show that *in vivo* sensory cortical networks do display low-exponent (see Supplementary Tables 1 & 2), power-law behavior for their response to varying stimuli intensity (as proxied by the number of spikes evoked in a target cell during stimulation). This is in line with predictions from critical dynamics, with the observed exponents closely matching the expected value (0.285) for a two-dimensional directed percolation system^28^. The expected exponent for the response scaling of a 3D network is 0.45, whereas higher dimensional systems would show an exponent of 0.5. Note that our observed ∼0.5 slope during baseline activity does not represent a response of the network: since neuronal firing during ongoing activity is correlated, that relationship is more indicative of autocorrelation in the network as a whole. Our measured scaling exponent thus emphasizes the intriguing finding that superficial layers 2/3, defined anatomically as an extended 2-dimensional sheet, functionally also operate more closely to a 2-dimensional critical network.

Our findings of maintained high Fano factor in evoked responses is in line with the expectation for a critical system, though, squares with the quenched variability demonstrated in response to sensory stimuli^36^. This difference may arise because single-neuron stimulation introduces a smaller perturbation than sensory stimulation, which in the awake state is dominated by inhibition^25^, affecting response variability. Early studies in anesthetized animals^18^ (e.g. in the visual cortex of cats) demonstrated that trial-by-trial variability in stimulus responses can be approximated as average response overlaid on a correlated, variable background. In the awake state, our observation of consistently high FF values for responses in the presence of scale-invariant parabolic avalanches indicates the need for a conceptual approach that bypasses the reliance on averaging. Avalanches emerge in superficial layers of cortex when animals recover from anesthesia^67,68^, exhibit high neuronal selectivity^29,69^ and spatiotemporal order^70–74^ characterized by power law statistics with exponents ≤ 2 that is, averages are not defined and variability is high. Yet, we demonstrate that decoding of novel spikes is similar across the fluctuation dominated avalanche regime.

Our decoding analysis provided important insights about the fine-grained nature of this communication of novelty. First, strongly decoding neurons were found in the immediate neighborhood of TCs supporting a spatial weighing in the response, in line with expectations from the local, synaptic connectivity statistics of pyramidal neurons. On the other hand, a large fraction of neurons with high decoding contribution did not fall into the PosR category or were located at long distances from the TC, supporting the contribution of indirect pathways.

Similarly, in our simulations the number of identified PosR greatly surpassed that of direct postsynaptic neurons of each TC, further suggesting the importance of indirect pathways to the coding of novel information. In fact, only about 16% of PosR had a direct connection from TCs. This number could be even higher for the brain, since the presence of inhibitory connections (absent in our simulations) could facilitate certain pathways in detriment of others, while also requiring stronger excitatory-to-excitatory connections in order to stay in a balanced propagation regime. This also de-emphasizes reliance on distinct, non-overlapping network connectivity.

The ability of PosR neurons to contribute significantly to the decoding of multiple TCs is in line with the spatial overlap of neuronal populations encoding different stimuli and their temporal separation. Our findings align with the notion that cortical networks are organized to maximize coding efficiency and redundancy, where overlapping populations can fluctuate in their response patterns to encode different stimuli^75,76^. The rapid decay found for the encoding of novel spikes further underscores this dynamic, ensuring that novel signals do not disrupt ongoing network activity but still convey novel information effectively within the constraints of the temporal and spatial coding architecture described for e.g. visual stimuli of the primary visual cortex in the awake mouse^77^.

Our results reveal the communication of single spikes within the scale-invariant, critical-state dynamics of cortical networks, where synchronized fluctuations are neither overly constrained nor entirely random. We propose this remarkable sensitivity of the mammalian cortex, to efficiently disseminate novel information from individual neurons, underscores the fundamental role of critical dynamics in cortical communication and computation.

## Materials and Methods

### Animal surgery

All procedures were approved by the NIH Animal Care and Use Committee (ACUC) and experiments followed the NIH *Guide for the Care and Use of Laboratory Animals*. Mice were obtained from Jackson labs, bred inhouse with C57BL/6 backgrounds (Jackson Laboratory) under a reversed 12:12 hr light/dark cycle. Chronic 2PI imaging in adult (>6 weeks) mice was enabled by using a head bar in combination with a cranial window placed centered at ∼2.5 mm from the midline (right hemisphere) and ∼1 mm rostral to the lambdoid suture and consisting of a stack of circular glass cover slips using established protocols^78^. Mice were injected with a 7:1 mixture of a viral construct to express jGCaMP7s^79^ and ChrimsonR^80^ in pyramidal neurons using the CaMKII promotor. A total of 2 – 3 injections of virus (100 – 400 nL; <1 µL in total; 10^13^ vg/mL; pGP-AAV-syn-jGCaMP7s-WPRE AAV9 & pAAV-CamKIIa-ChrimsonR-mScarlet-KV2.1, Addgene) were administered into the right hemisphere. Chronic 2PI started after >2 weeks in identified V1 at a depth of ∼100–200 μm.

### Identification of V1 maps

Retinotopic maps of V1 and higher visual areas (HVAs) were generated for all mice prior to recording using published protocols^81,82^. Briefly, awake, head-fixed mice faced with their left eye a 19” LCD monitor placed at 10 cm distance and tilted 30° towards the mouse’s midline. Using Psychophysics toolbox^83^, contrast-reversing, spherically corrected checkerboard bars were drifted across the screen vertically (altitude) and horizontally (azimuth) for each of the four directions (30 repeats per direction). Simultaneous wide-field imaging (Quantalux, Thorlabs) captured jGCaMP7s fluorescence, which was averaged for each direction. Altitude and azimuth phase maps were calculated by phase-wrapping the first harmonics of the 1D Fourier transform for each of the four averages and subsequently subtracting the maps of the opposite directions^82^. Sign maps were generated by taking the sine of the angle between the gradients in the altitude and azimuth maps and processed^81^. Borders were drawn around visual area patches and overlaid onto anatomical reference images to identify V1.

### Holographic stimulation of single neurons

Holographic stimulation^84^ was done by manually selecting individual pyramidal neurons as target from the two-photon image of the field of view in both red and green channels. Neurons that express the opsin (ChrimsonR, with mScarlet tag) were selected for stimulation. Based on the selected targets, a map of top-hat patterns with diameter of 10 µm was used to generate the hologram on the Spatial Light Modulator (SLM, Meadowlark, LCoS high resolution 1920×1152 phase modulator). Light from the laser/OPA system (Carbide/Orpheus-F – Light Conversion, λ = 1064 nm, repetition rate = 600 kHz, pulse width <100 fs) was expanded (4x) to fill the area of the SLM (8 mm × 8 mm). The hologram on the pupil plane was shrunk to fit the size of the Galvo-Galvo scanners (∼3 mm diameter) and form the desired pattern of top hats after the objective lens (Nikon 16X, working distance = 3mm and effective focal length = 12.5 mm) on the selected targets. Each target neuron was continuously stimulated by the top-hat beam shape for 100 ms for a total of ∼100—150 trials per experiment, with an interval of 2 s (10 s in some recordings) between trials. Power at target neurons ranged from ∼2.5 – 10 mW. It is important to highlight that no artifact from this stimulation was detectable on our imaging with our combination of GECI/opsin, stimulation wavelength and the low power required to stimulate single targets (see Supplementary Fig. 4).

### 2PI imaging, pre-processing pipeline and meta data collection

For standard 2PI, images were acquired by a scanning microscope (Bergamo II series, B248, Thorlabs Inc.) coupled to a pulsed femtosecond Ti:Sapphire 2-photon laser with dispersion compensation (Chameleon Discovery NX, Coherent Inc.). The microscope was controlled by ThorImageLS and ThorSync software (Thorlabs Inc.). The wavelength was tuned to 940 nm to excite jGCaMP7s. Signals were collected through a 16× 0.8 NA microscope objective (Nikon). Emitted photons were collected through a 525/50 nm band filter using GaAsP photomultiplier tubes. The field of view was ∼450×450 μm. Imaging frames of 512×512 pixels were acquired at 45.527 Hz by bidirectional scanning of a 12 kHz Galvo-resonant scanner. Beam turnarounds at the edges of the image were blanked with a Pockels cell. The average power for imaging was <70 mW, measured at the sample.

The obtained tif-movies in uint16 format were rigid motion-corrected via the python-based software package ‘*suite2p*’^85^. Registered images were further denoised using machine-learning based, deep interpolation^86^ (see below) and then semi-automatically processed by suite2p for ROI selection (we performed visual curation of the automatically-selected ROIs to ensure only neurons were included in the analysis; this is done so processes, such as the dendrites identified in Fig. 1b, are not part of the analysis and therefore cannot be classified as significant responders or contribute spikes to response curves or other measurements) and fluorescence signal extraction. For each labeled neuron, raw soma and neuropil fluorescence signals over time were extracted for each ROI. Spiking probabilities were obtained from neuropil-corrected fluorescence traces (F_corrected_ = F_ROI_ – 0.7*F_neuropil_) via MLspike (https://github.com/MLspike), utilizing its autocalibration feature to obtain unitary spike event amplitude, decay time, and channel noise for individual ROIs.

#### Deep-interpolation

Deep-interpolation^86^ (Deep-IP; https://github.com/AllenInstitute/deepinterpolation) removes independent noise by using local spatiotemporal data across a noisy image stack of N_pre_ + N_post_ frames to predict, or interpolate, pixel intensity values throughout a single withheld central frame. The deep neural network is a nonlinear interpolation model based on a UNet inspired encoder-decoder architecture with 2D convolutional layers where training and validation are performed on noisy images without the need for ground truth data. As described previously in detail^29^, after rigid motion correction, individual denoised frames were obtained by streaming one 60-frame (N_pre_ = N_post_ = 30 frames) registered, image stack through the provided Ai-93 pretrained model^86^ for each frame to be interpolated. At an imaging rate of ∼45 Hz, these 60 frames correspond to a combined ∼1.3 seconds of data surrounding the frame to be interpolated. This process did not alter qualitatively our main results (see Fig. 4f & Supplementary Fig. 8).

### Postprocessing pipeline

Our postprocessing pipelines were custom-written in Matlab (Mathworks) and Python (www.python.org). Some routines utilized NumPy (https://numpy.org/) and Matplotlib (https://matplotlib.org/).

#### Stimulus response measurement

Unless stated otherwise, spike count over a window of 6 imaging frames (∼132 ms) starting at the stimulation onset was used as a measure of the response to holographic stimulus of each target cell for all cells. That window was chosen because it includes the full stimulus duration (100 ms) and one extra frame after the stimulus offset, for which the increased activity in TC can still be observed (see Fig. 1f). This was compared to the baseline count, defined as a 6-frame window starting ∼264 ms before stimulus onset. Due to difficulties and limitations of spike deconvolution of calcium traces, some parts of the time series resulted in artificially high spike rates. To avoid artificial increase of variability measures due to these artifacts, we introduced an outlier removal procedure in which trials with spike count above 10 standard deviations over the mean are removed from the analysis. This procedure resulted in <0.1% trials removed, across all neurons for all experiments.

#### Target selection and exclusion

Up to 25 target cells were visually selected based on expression and location in the field of view (cells close to the edges were avoided) for each experiment. Not all targets were responsive to stimulation. Targets in which fewer than 20% of the trials evoked a spike count above the 91% percentile of its baseline count were excluded from the analysis. This procedure resulted in 54 target cells across 16 experiments from 6 mice being considered out of 193 targets attempted across 33 experiments from 11 mice. We estimate an average of 46.3 ± 17.9 ChrimsonR-labeled cells per field of view, which translates to an average density of labeled cells of ∼2.3×10^-2^ cells per 100 µm^2^. This sparse labeling facilitates the holographic stimulation of individual targets (see examples at Supplementary Fig. 1).

#### Definition of significant responders

Cells with significant response to each target’s stimulation were defined as those whose spike count distribution was significantly different from its baseline count distribution. Specifically, we used a z-test to compare the stimulus response spike count to the average and standard deviation of the baseline count for each non-target cell. A cell was defined as a positive responder if the z statistic is above the highest α = 2.5% (unless stated otherwise) of the distribution and as a negative responder if the z statistic is below the lowest α = 2.5%.

Otherwise, the cell is classified as a non-significant responder. We have also corrected for false-discovery rate (FDR), by employing the matlab function mafdr ^34,35^, with an FDR of 2.5%. After FDR correction, the number of positive responders increased in relation to the one obtained at α = 2.5%, whereas the number of negative responders dropped to nearly zero. It is important to highlight that, given a low baseline activity rate for most cells, with an average count in the relevant window for our analysis near zero, it is technically challenging to define negative responders in this scenario using spike data. Other groups have shown suppressed neurons upon select pyramidal neuron stimulation using fluorescent traces, for which detection of significant decreases during stimulation might be easier^7,87^.

This procedure is done for each target stimulated in an experiment separately. Therefore, in experiments where multiple targets were selected, each non-target cell is tested for significance of its response to each target’s stimulation, possibly being a significant responder for some targets and non-significant responder to others (see Fig. 4d).

#### Fano Factor calculation and significance

Fano Factor was calculated as usual, for each cell separately, by dividing the variance of their spike count by the mean of their spike count across trials. To correct for changes in firing rate across conditions^37^, we introduced a procedure in which the window used to measure the spike count during stimulation varied from 1 to 9 frames (∼22 ms to 198 ms), starting at the onset. The window length was determined by minimizing the normalized difference between the mean baseline and stimulus counts. Cells for which this difference was above 20% were excluded from the analysis. This procedure resulted in less than 30% of cells removed, on average, across the non-target cells and an average of 65% of targets removed (minimum of 13% or 7 targets considered). Values were reported as mean ± error over the mean across cells, unless stated otherwise. Significance of Fano Factor changes across conditions was obtained by employing the Wilcoxon rank sum test at a significance level of 0.05 (p-values reported in Results section and figure legends).

#### Trial separation and controls

To control for different conditions affecting the variability, the Fano Factor was recalculated and compared using a subset of trials. To control for target cell response variability affecting non-target cells, we separated trials into low target count trials and high target count trials. We defined low count trials as those for which the target cell produced a spike count during stimulation that was above the 91^st^ percentile of the baseline count distribution and up to twice that number of spikes, whereas high count trials were any trial with count above that limit. To control for the influence of ongoing fluctuation of the network activity we separated trials into low and high population baseline count, defined as trials for which the population count during baseline was lower than the 25^th^ percentile of its distribution or higher than 75^th^ percentile, respectively. To control for temporal influence of the responses to the stimulus, we defined two consecutive 6-frame windows, centered at the peak of the target’s average response to the stimulus (2—3 frames after onset), as early and late response windows.

#### Correlation analysis

Spike counts across trials were compared by computing correlations across time (autocorrelation, calculated using the spike counts over pairs of windows separated by a temporal lag of 132 ms) and across cells (comparing spike counts of targets *vs* non-targets). Values are reported as mean ± error over the mean. Significance of correlation changes across conditions was obtained by employing the Wilcoxon rank sum test at a significance level of 0.05 (p-values reported in Results section and figure legends).

### Decoding Analysis

#### Classification model selection, performance metrics, and data preprocessing

In our prediction and classification scheme, we used target neuron ensembles as “classes” and the activity of the responding/non-target neurons as the “features”. As described in Stimulus response measurement section, we use spike count over a window of 6 imaging frames as a feature value for each neuron. For the temporal analysis, this window is offset relative to stimulus onset, as described below. In a preliminary analysis, we compared several machine learning strategies and found that the ensemble methods of boosting (XGBoost^38^) and bagging (scikit-learn^88^ Random Forest classifier) performed best to several other common classification schemes such as Logistic Regression, Naive Bayes, Neural Networks, and Support Vector Machines. These two approaches yielded the highest accuracy and were then applied to all of our analyses presented in this work. In all cases we split the data samples randomly so that 80% was used for training and 20% for testing. We repeat this split 100 times and from these report the mean and standard errors of our metrics. As metrics we use overall accuracy, per-class recall and precision, as well as F1-scores (harmonic mean of recall and precision). For the results in Fig. 4f and Supplementary Fig. 10d we use all data and all trials, but for others we performed additional pre-processing before running classification, as detailed below.

Since not all trials were equally successful in eliciting spikes from target cells, for all other decoding analysis we selectively filtered trials and targets, with a criterion that the target neuron had to produce at least one spike across any of the six frames after the onset of stimulation. We then selected valid targets as those that spiked on at least half of their trials. Thus, non-significantly driven TC (based on conservative response spike count) excluded some TC in the spike count analysis which nevertheless had significantly F1-score in the decoding task (gray TC in Supplementary Fig. 10a). To characterize feature importances and interpret our classification schemes we used Shapley analysis^89^, as described below.

#### Decoding from baseline

To examine whether TCs are decodable during spontaneous activity, we applied a similar analysis to the background activity in the periods between stimulation trials. We segmented this background activity using the same bin size (6 frames) as in the stimulation condition and identified pseudo-trials in which only one of the previously used TCs spiked within a bin. To avoid potential artifacts introduced by Deep Interpolation, we excluded windows immediately preceding or following stimulation. For each of these background spike datasets, we restricted the analysis to the subset of TCs used during decoding in the stimulation condition and downsampled the data to match the sample distribution of the stimulation trials (Fig. 4b, *right*).

#### Radius Analysis

For each target neuron, we implemented an exclusion zone based on the distance from the target neuron’s soma, *r*. We progressively increased *r*, and the neurons/features falling within this distance were systematically eliminated from the feature matrix, enabling quantification of the relationship between F1-score degradation and exclusion zone radius for individual target neurons. We computed the probability distribution of any feature neuron being excluded, as a function of distance, P*_ex_*(r). Specifically, we aggregated the frequency of neuron eliminations at each radius across all datasets and normalized by the total number of exclusions.

The spatial dependency of predictive performance was characterized by analyzing the differential changes in F1-score across the range of exclusion radii. For each target neuron across all datasets, we identified the radius at which the decrease in F1-score first exceeded three standard deviations below the mean rate of change. These radii were averaged across datasets and smoothed using a Gaussian kernel (σ = 1) to obtain an estimate of the spatial scale of predictive influence.

#### Neuron Drop Analysis and Positive Responder Probability

To evaluate the distributed nature of neural information and assess the robustness of our decoding framework, we conducted an iterative feature elimination analysis for each target neuron across all datasets. For each iteration, including initial model fitting, we use Shapley values (employed using SHAP package: SHapley Additive exPlanations^90^) to quantitatively rank feature neurons based on their predictive contribution for a given target neuron. Through sequential elimination of the highest-ranked feature neurons and subsequent model retraining, we systematically assessed how the removal of informative neurons impacted decoding performance. This iterative process of SHAP-guided feature elimination and model reconstruction was repeated until a single neuron remained. At each iteration we recorded F1-score, precision, and overall accuracy and the entire procedure was replicated one hundred times, utilizing different out-of-bag random selection of training and testing sets.

To characterize the relationship between feature importance and significant responders, we quantified the probability that eliminated neurons exhibited significant responses. For each target neuron, we constructed a binary vector corresponding to the sequential elimination order, where entries were assigned values of 1 or 0 based on whether the eliminated neuron was a significant responder. These binary sequences were averaged across all iterations of the elimination procedure to obtain target-specific probabilities and then averaged across all target neurons.

#### Temporal profiles of the prediction accuracy

To further characterize our holographic stimulation data we explored the accuracy of our prediction scheme as a function of time. This procedure allowed us to study how long the information about the origin of stimulation persists in network. Note that the prediction scheme is a more stringent requirement than studying only the causal effects of stimulation, since even though the activity after the stimulation can reverberate in a network for a long time, identifying the node that initiated it is more challenging. Furthermore, for the temporal analysis we disabled our filtering of inactive targets/trials, utilizing all data. This simplifies the interpretation of these results, with an expected overall decrease in accuracy compared to the other decoding analyses.

We assessed the temporal course of the prediction accuracy across datasets, each with different levels of responsiveness of the targets. We varied the size of our response window (1 or 6 frames) and then moved it in steps of 1. The main results are present in Figure 4f (single-frame analysis) & Supplementary Fig. 10d (six-frame analysis). For the six-frame analysis, the fading “pink” rectangles indicate the amount of data used from the stimulation region (100 ms) window, so that the fully white regions use data strictly before or after stimulation is over. Note that the bins and the stimulation region are not perfectly aligned, as the trigger for the stimulation was independent from the image scanning raster. We define the reference bin 0 as the first bin which fully overlaps with the stimulation.

### Avalanche analysis

#### Continuous epochs of suprathreshold population activity

Continuous periods of population activity were identified by applying a threshold *Θ* on the population activity *p*(*t*), the sum of the spikes from all neurons at a given time *t*, such that:

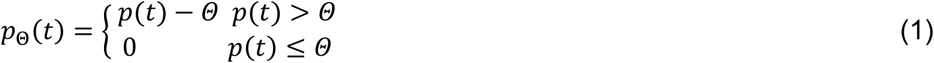

This procedure is known as soft-thresholding and was employed for all analysis unless otherwise stated. For a given recording *p*(*t*) and coarse-graining value *k*, the dependence of the number *N* of epochs on the threshold *Θ*, N(*Θ*) was obtained for a range of thresholds *Θ* ɛ [*Θ*_1_, *Θ*_2_] such that *Θ*_1_ was low enough that it removed no population activity from the time course and *Θ*_2_ was high enough that it would remove all population activity from the time course. The function N(*Θ*) was typically well-approximated by a log-normal distribution and a corresponding fit yielded shape parameters *μ* and *σ*. The threshold used in the analysis for all recordings and coarse-graining factors was chosen such that *Θ* = μ and estimated for each *k*. See Capek & Ribeiro et al.^29^ for more details.

#### Temporal coarse-graining

A temporal coarse-graining operation was applied to the thresholded population activity *p*_*Θ*_(*t*). For a given temporal coarse-graining factor *k* an ensemble of *K* different coarse-grained time series 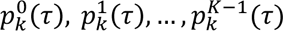 was arrived at through the following method:

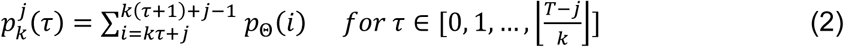

For each time series 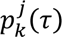, epochs were extracted by finding pairs (*τ*_1_, *τ*_2_) such that 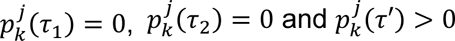 for all *τ*^′^ ∈ [*τ*_1_+ 1,…, *τ*_2_− 1]. The size of the epoch is given by 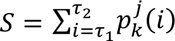 and its corresponding duration given by *τ*_2_ – *τ*_1_ − 1. For a given ensemble of coarse-grained time series, all epochs were pooled.

#### Scaling curve fit

For more precise evaluation of the scaling curve, mean avalanche size *vs* duration, we introduced the following fitting function:

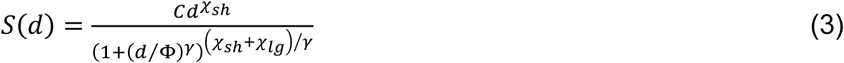

This function is a double power law with initial slope *χ*_sh_, transitioning to a second slope *χ*_lg_ at around the point *d* = Φ. The parameter γ controls how abruptly that transition happens and has been fixed at 4 for all the curves presented. The other parameters were free to adjust to the data, and all fits were performed in log-space, i.e., the log(*S*(log(*d*)) was fit to the log of data (taking the log of both average sizes as well as durations).

### Simulations

We created a 300×300 two-dimensional lattice of excitable cellular automata^91^. We defined the spacing of the lattice as 12 μm, to mimic the high density of cells in brain tissue. We connected cells by a probability function that changes as function of the distance between cell pairs, with a short (gaussian, 100 μm characteristic decay) and a long (exponential, 290 μm characteristic decay) component: 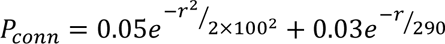. At each time step (set at 2 ms to mimic average spike transmission times in the brain), each cell can be in one of three states: 1) resting, 2) active, or 3) refractory. A resting cell can become active by either receiving an external Poisson drive (with probability P_poiss_) or by an active presynaptic neighbor (with probability P_trans_). We set 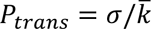, where 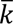 is the average number of connections per cell (∼43 with the numbers described above) and σ is the branching parameter, a variable we can adjust to move the system in and out of criticality and is expected to be slightly above 1 at the critical point (to compensate for energy loss/collisions in spike propagation in the network – ∼1.0066 on our simulations). We set *P*_*poiss*_ (*σ*), so that the average amount of activity during baseline for each σ value matches the average baseline activity in the experiments (see Fig. 6d).

We then selected for each “experiment” (10 total), a non-overlapping set of 10 cells in the center of the network (inner 38×38 cells) to be the target of extra external drive (as in the mice experiments, for 100 ms or 50 time steps, with 2 s inter-trial interval), which was set so the average activity of each target in the simulations during stimulation matches the average activity of targets during stimulation in the experimental data. All analyzes are then performed exactly as done for the experimental data, but only using the inner 38×38 cells, to avoid border effects (periodic boundary conditions were used to simplify the simulations).

#### Output Strength

To evaluate the relationship between the structure of the network and each target neuron’s ability to spread information regarding their firing, we introduced the measure of output strength: 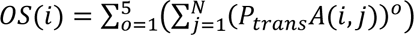, where *A* is the network’s adjacency matrix (*A(i,j)* = 1 when there is a connection from neuron *i* to neuron *j*, and 0 otherwise), *N* is the number of neurons and *P_trans_* is the spike transmission probability. By taking *P_trans_ A* to the power *o*, we evaluate the likelihood for a spike to transmit from *i* to *j* via an *o*-order path (e.g., order 1 would be a direct connection, order 2 would be a two-step connection – neuron *i*, to neuron *k*, to neuron *j*).

Therefore, after summing over all other neurons, *OS(i)* estimates the number of spikes generated in the network for each spike neuron *i* produced, excluding dynamics (collisions, refractory periods, etc.) and only considering the network’s structure.

## Data availability

The preprocessed imaging data used in this study are available in the general repository Zenodo using the following access DOI:TBD. The source data for all figures and supplemental figures in this study are provided for this paper.

## Code availability

Computer code used in this study is available at the following github link: TBD.

## Acknowledgements

We thank Craig C. Stewart and members of the Plenz lab for help with animal surgery, animal care. This research was supported by the Division of the Intramural Research Program (DIRP) of the National Institute of Mental Health (NIMH), USA, ZIAMH002797, ZIAMH002971, and the BRAIN initiative Grant U19 NS107464-01.

This research was supported by the Intramural Research Program of the National Institutes of Health (NIH), USA, ZIAMH002797, ZIAMH002971, and the BRAIN initiative Grant U19 NS107464-01. This research utilized the supercomputing resources of the National Institutes of Health (NIH, USA; Biowulf, http://hpc.nih.gov).

The contributions of the authors were made as part of their official duties as NIH federal employees, are in compliance with agency policy requirements, and are considered Works of the United States Government. However, the findings and conclusions presented in this paper are those of the author(s) and do not necessarily reflect the views of the NIH or the U.S. Department of Health and Human Services.

## Author Contributions

TLR and DP conceived and planned the study; TLR, BG, and VS performed experiments; AV took the lead in holographic setup design. TLR took the lead in data analysis with support from BG and VS. SP and RS took the lead on machine learning based decoding of experimental data. All authors contributed to the analyses. TLR and DP wrote the manuscript.

## Supplementary Figures

**Supplementary Fig. 1.**
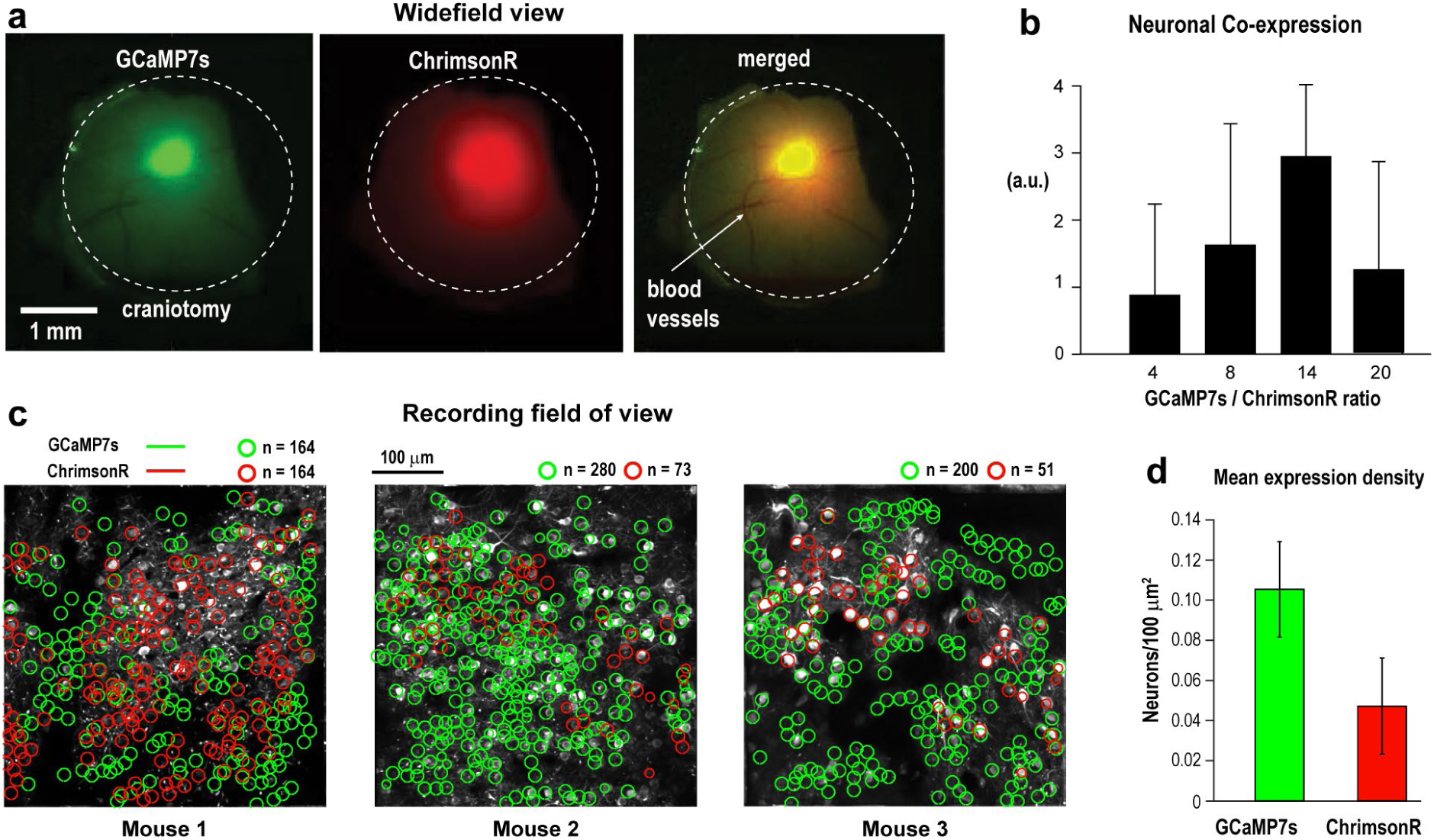
**a** Widefield image showing GCaMP7s (*green*) and ChrimsonR (*red*) expression, with the merged image indicating co-expression (single mouse). **b** Optimal co-expression achieved with a virus titer ratio of 14:1 (n = 23 mice). **c** Examples of individual neurons with significant GCaMP7s (*green*) and ChrimsonR (*red*) expression from 3 mice. **d** Mean expression density showing twice as many neurons positive for GCaMP7s compared to ChrimsonR (from c).

**Supplementary Fig. 2.**
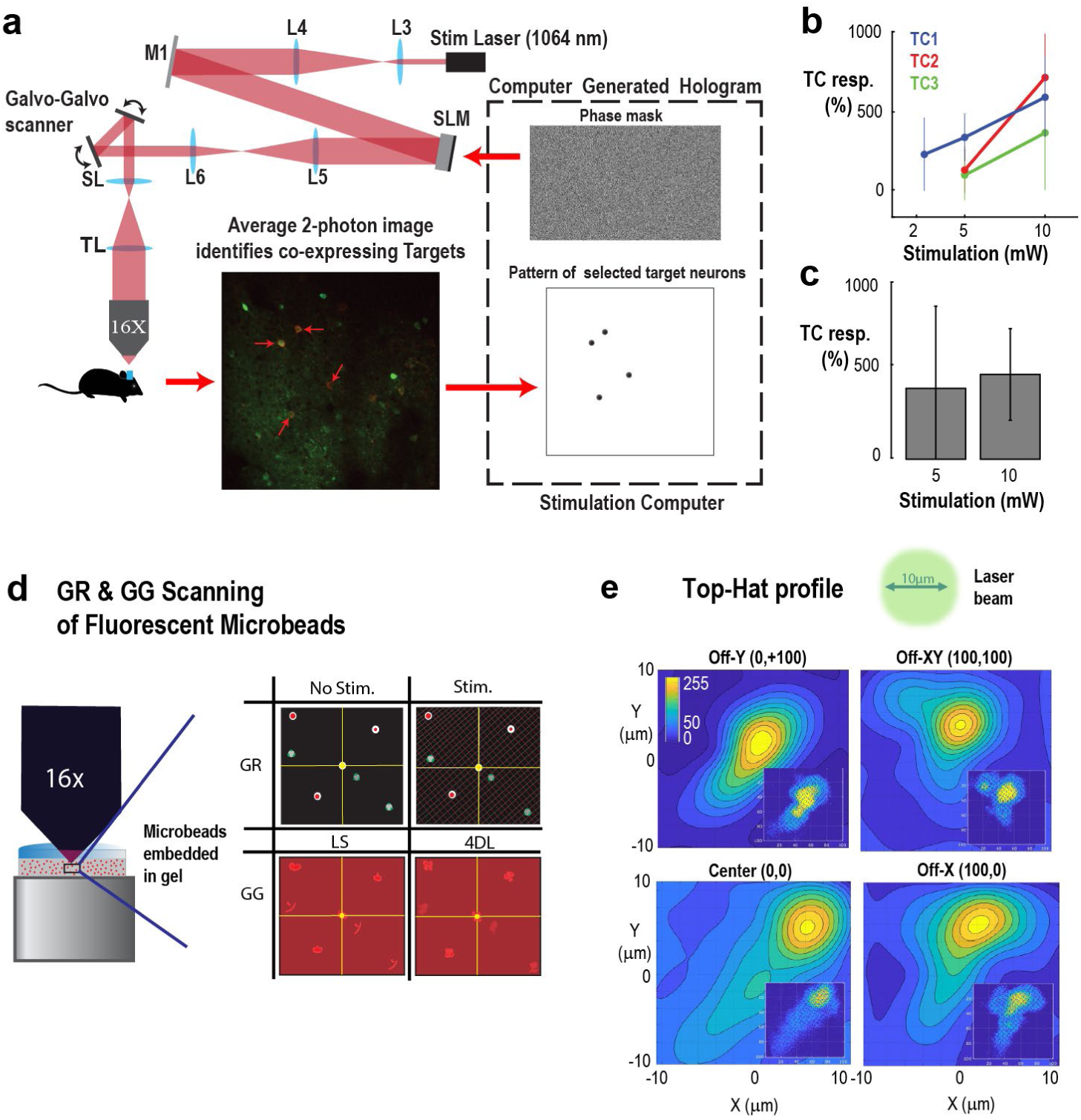
Schematic design and evaluation of the stimulation path using spatial light modulation. **a** Laser light (Light Conversion, 1064 nm, 600 kHz, <100 fs) is expanded to fill the area of the spatial light modulator (SLM; Meadowlark, 1920×1152 resolution). The hologram is resized to fit the Galvo-Galvo scanners (∼3 mm diameter), which guide the stimulation beam in the focal image plane (Nikon 16X, 3 mm working distance, 12.5 mm effective focal length). Target cells (TC) expressing both GCaMP7s and ChrimsonR (opsin with tdTomato tag) are selected from the mean 2PI image of the field of view from both red and green channels for stimulation and simultaneous estimation of the evoked spike count. Map of TC informs a hologram of top-hat patterns with diameter of 10 µm for the SLM. Optical power was adjusted to 5—10 mW per TC, depending on the depth and expression level. **b** Spiking above base line increases with stimulation power (N = 3 TC; 100 ms stimulation duration). **c** No difference in evoked spike count for 5 and 10 mW stimulation across all TC. **d** Schematics of the stimulation xy-profile reconstruction using fluorescence microbeads. Galvo-Galvo (GG) controlled stimulation of gel-embedded microbeads (3 µm diameter) and simultaneous Galvo-Resonance (GR) 2PI of elicited response. *Circles*: Individual microbeads before and during stimulation. **e** Experimentally reconstructed and smoothed xy-profile of a flat-hat stimulation (10 µm diameter; 5 mW) at 4 different positions within the field of view at center (0,0), off-x (+100 µm, 0), off-y (0, +100 µm), and off-xy (+100 µm, +100 µm). *Inset*: Non-smoothed reconstruction.

**Supplementary Fig. 3.**
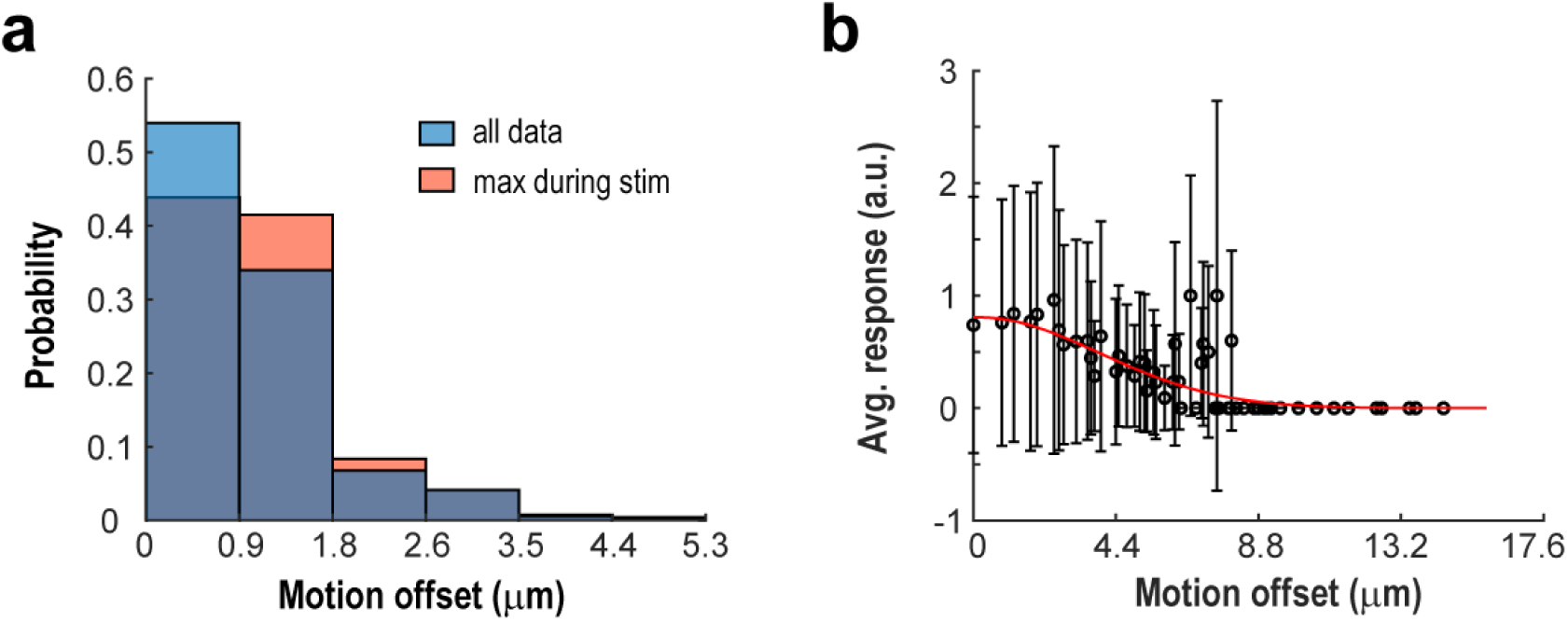
Analysis of motion correction from processed recordings reveals low amount of motion artifact affecting stimulation. **a** Distribution of motion offsets observed on each frame over all recordings. *Blue*: all data. *Orange*: maximum displacement observed during stimulation for each trial. **b** Spatial profile of stimulation response from targets measured using motion offset in trials during stimulation. Note that responses drop to approximately zero in frames with an offset above ∼4.4 μm.

**Supplementary Fig. 4.**
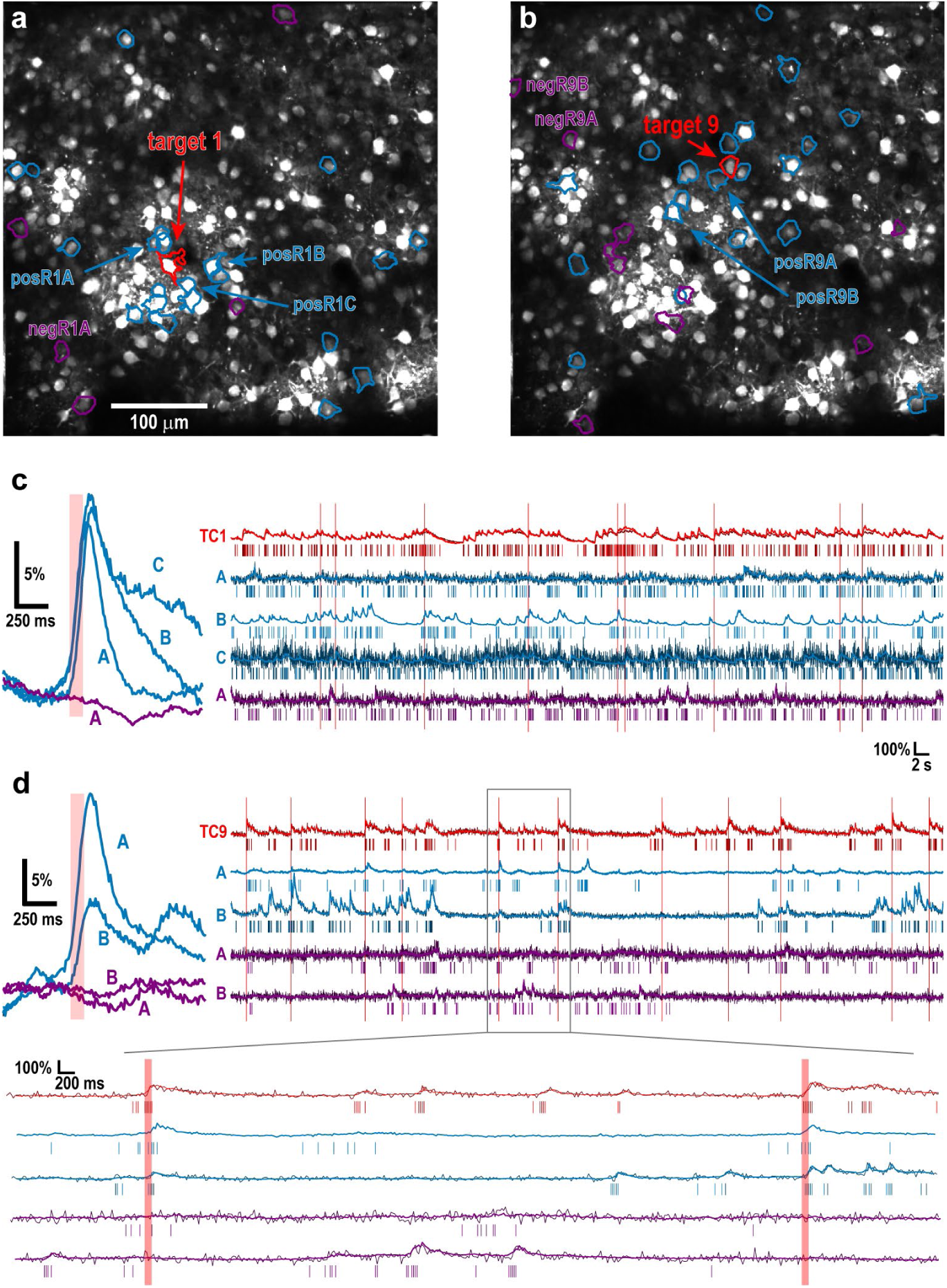
Example recording with activity from two example targets and some of their significant responders highlighted. **a** Example field of view with target 1 highlighted in *red* and its positive (*blue*) and negative (*purple*) responders indicated. **b** Same recording as in a, but now highlighting target 9 and its significant responders. c *Left*: time average fluorescent response to stimulation for the significant responders labeled in a. *Right*: example raw (*dark*) and denoised (*light*) fluorescent traces (*lines*) and deconvolved spikes (*bars*) for the labeled neurons in a. d *Left*: time average fluorescent response to stimulation for the significant responders labeled in b. *Right*: example raw (*dark*) and denoised (*light*) fluorescent traces (*lines*) and deconvolved spikes (*bars*) for the labeled neurons in b. *Bottom*: zoom in from the right panel above.

**Supplementary Fig. 5.**
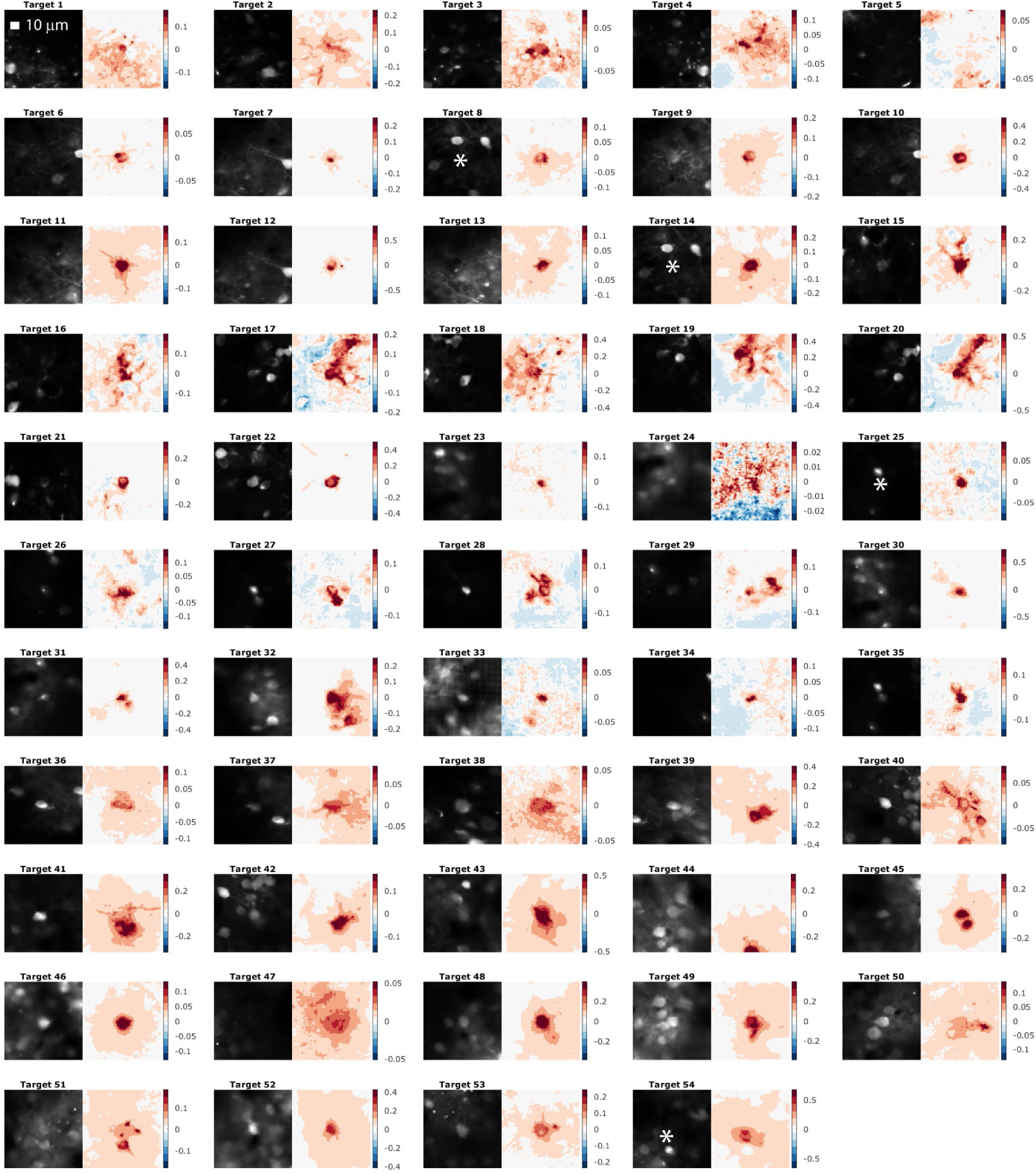
Summary of isolating individual pyramidal neurons in layer 2/3 of the primary visual cortex during rest using holographic stimulation and 2PI. For each target cell (TC), mean luminance after motion correction and denoising (*left*) and the ΔF/F in response to stimulation in a 100×100-pixel area (∼88×88 µm) centered on the TC (*right*) are shown. *Asterisks* are visual guides for TC position.

**Supplementary Fig. 6.**
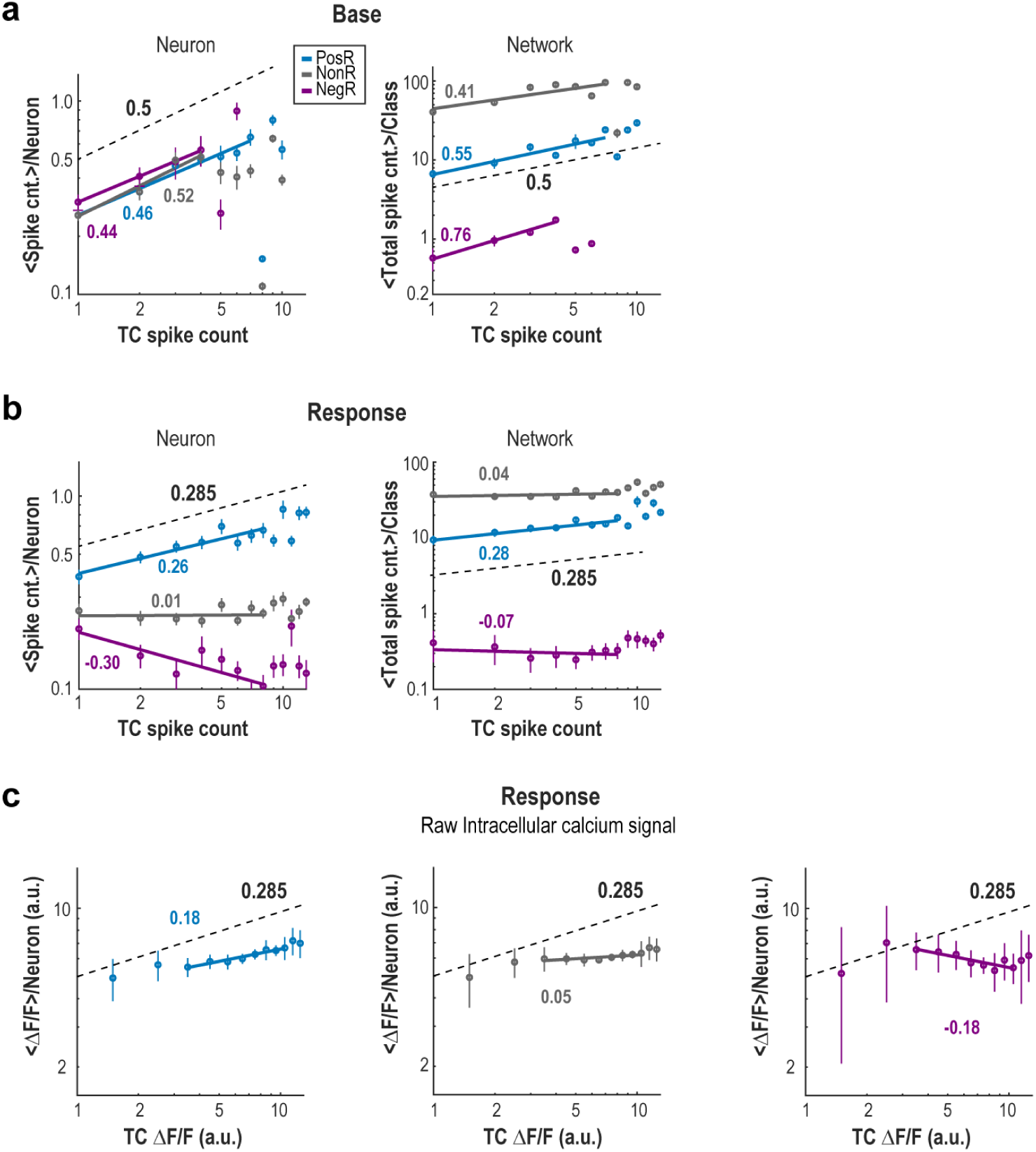
Spike count in all different subpopulations scale with spike count of target cells during baseline activity, but only Positive Responders scale during stimulation. **a** Mean spike count on PosR (*blue*), NonR (*grey*) and NegR (*purple*) as function of spike count of TC, in log-log, during baseline. Power law fits are indicated by solid lines, with obtained exponents shown. *Left*: mean calculated per neuron. *Right*: mean calculated after summing over the population. **b** Same as in a, but during holographic stimulation. **c** Response scaling calculated using the fluorescent traces (ΔF/F).

**Supplementary Fig. 7.**
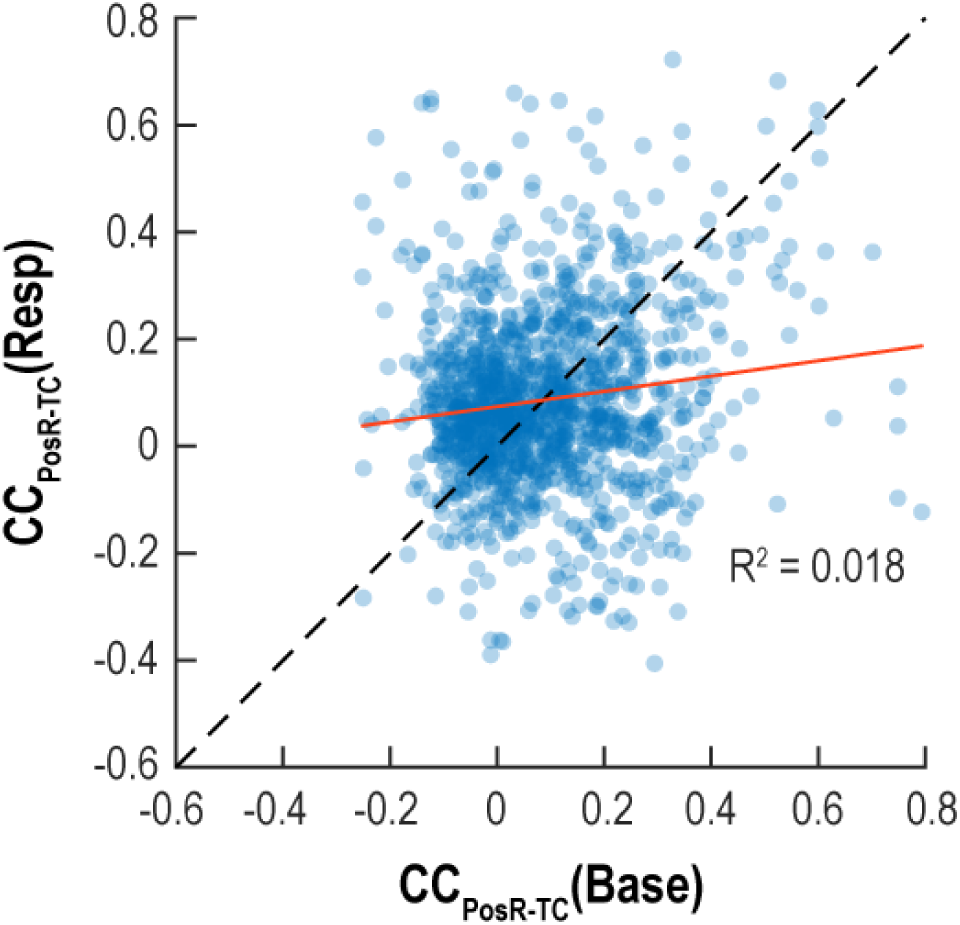
Correlations between PosR and TC change during stimulation in comparison with baseline. For each PosR-TC pair, their spike count correlation during stimulation is plotted against baseline. Note the shift towards higher values during Resp compared to Base (cp. Fig. 2c). A linear regression (*orange line*) shows that these measures are very weakly related (R^2^ = 0.018; linear regression; p < 10^-5^, for the *t*-statistic of the two-sided hypothesis test), indicating that the stimulation of TCs engages both existing but also new networks.

**Supplementary Fig. 8.**
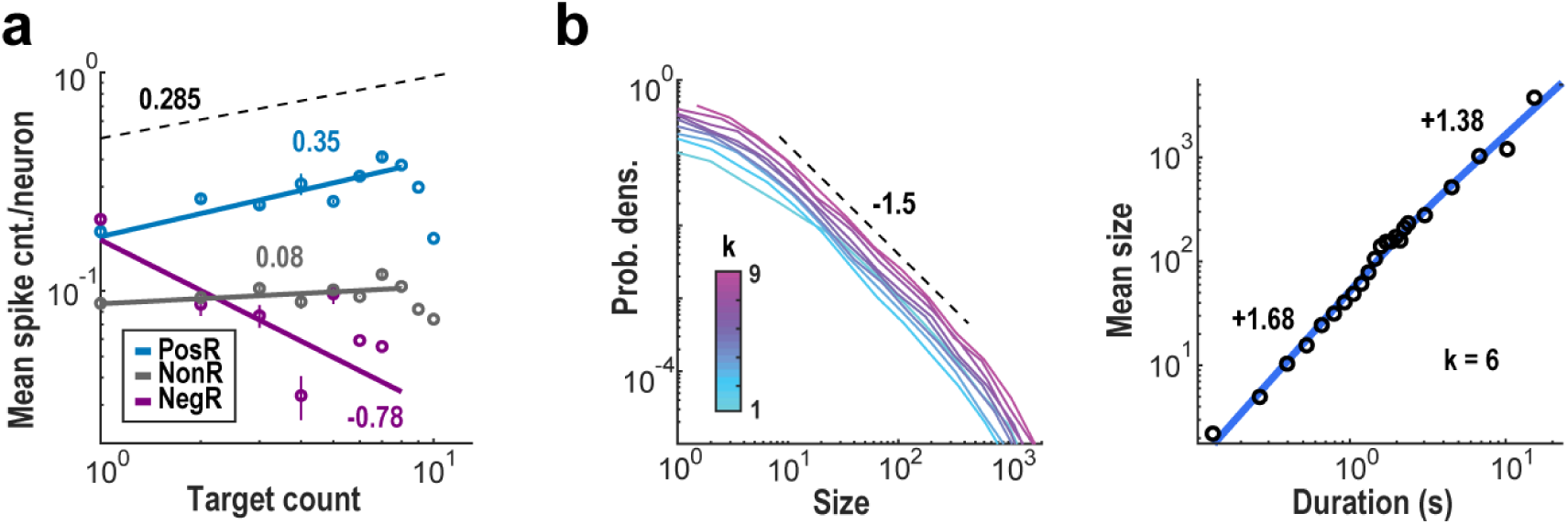
Response scaling and avalanche analysis are qualitatively unchanged without denoising from Deep Interpolation. **a** Average response from PosR (*blue*), NonR (*grey*) and NegR (*purple*) as function of TC spike count during stimulation, for datasets processed without denoising (no Deep Interpolation). **b** Avalanche analysis for datasets processed without denoising. *Left*: Distribution of avalanche sizes for different coarse graining levels *k* (*color bar*). *Right*: Mean avalanche size as function of avalanche duration at *k* = 6 showing an exponent below 2 for the short avalanches regime, in line with previous results obtained from noisy datasets (Ribeiro, Capek et al., 2023).

**Supplementary Fig. 9.**
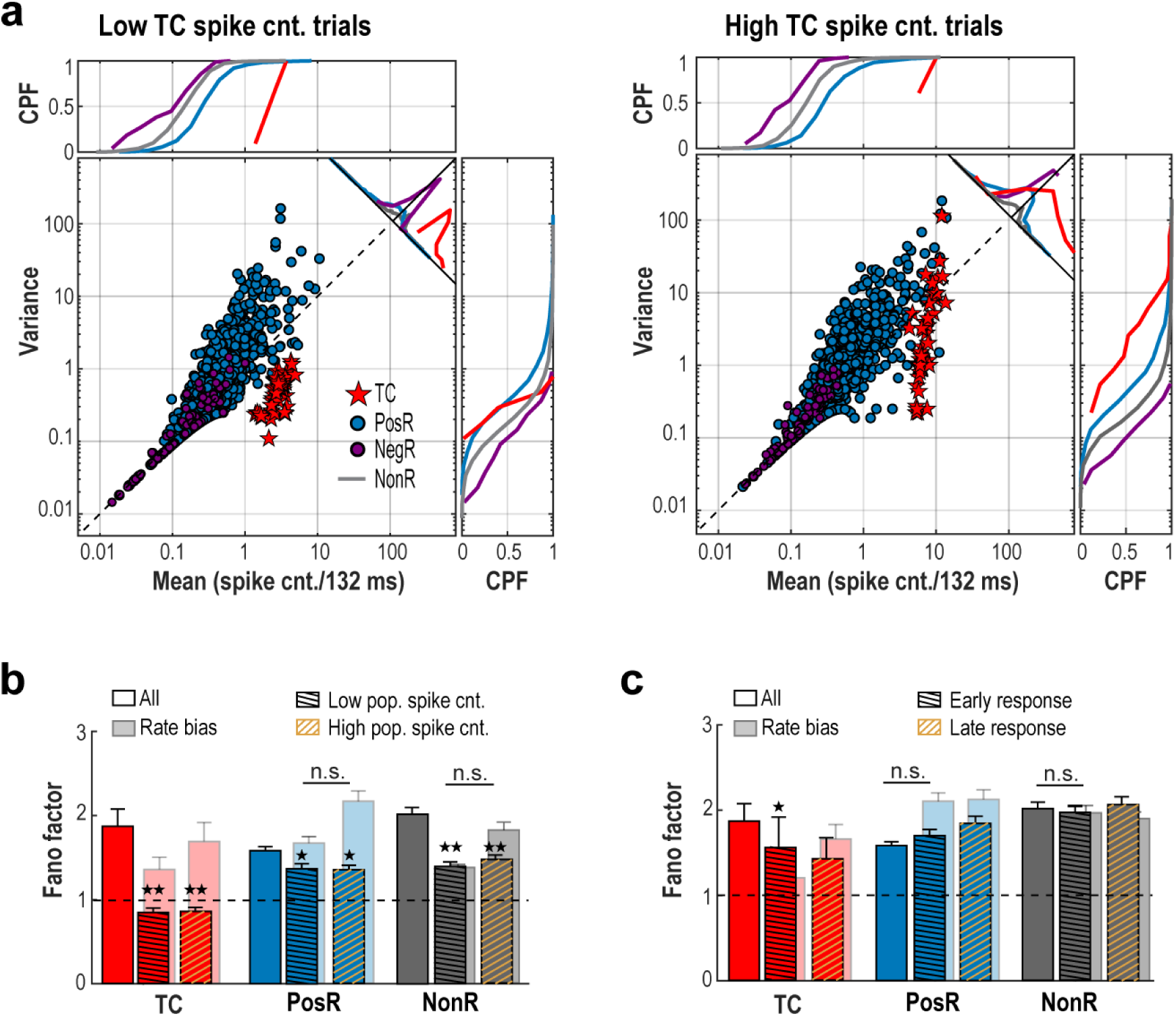
High response variability in PosR and NonR does not originate from variability in evoked spikes of TC. a Summary statistics of mean TC trials separated by low and high target counts. Note the remaining high variance of PosR despite the large drop in TC variance. No change in Fano Factor (FF) when subdividing into TC trials with high or low preceding population spike count (b) or early (first half of the response window) vs. late (second half of the response window) spiking responses (c). When comparing FF (b, c), 1 & 2 stars indicate p < 0.05 & 10^-3^, respectively, using Wilcoxon rank sum test.

**Supplementary Fig. 10.**
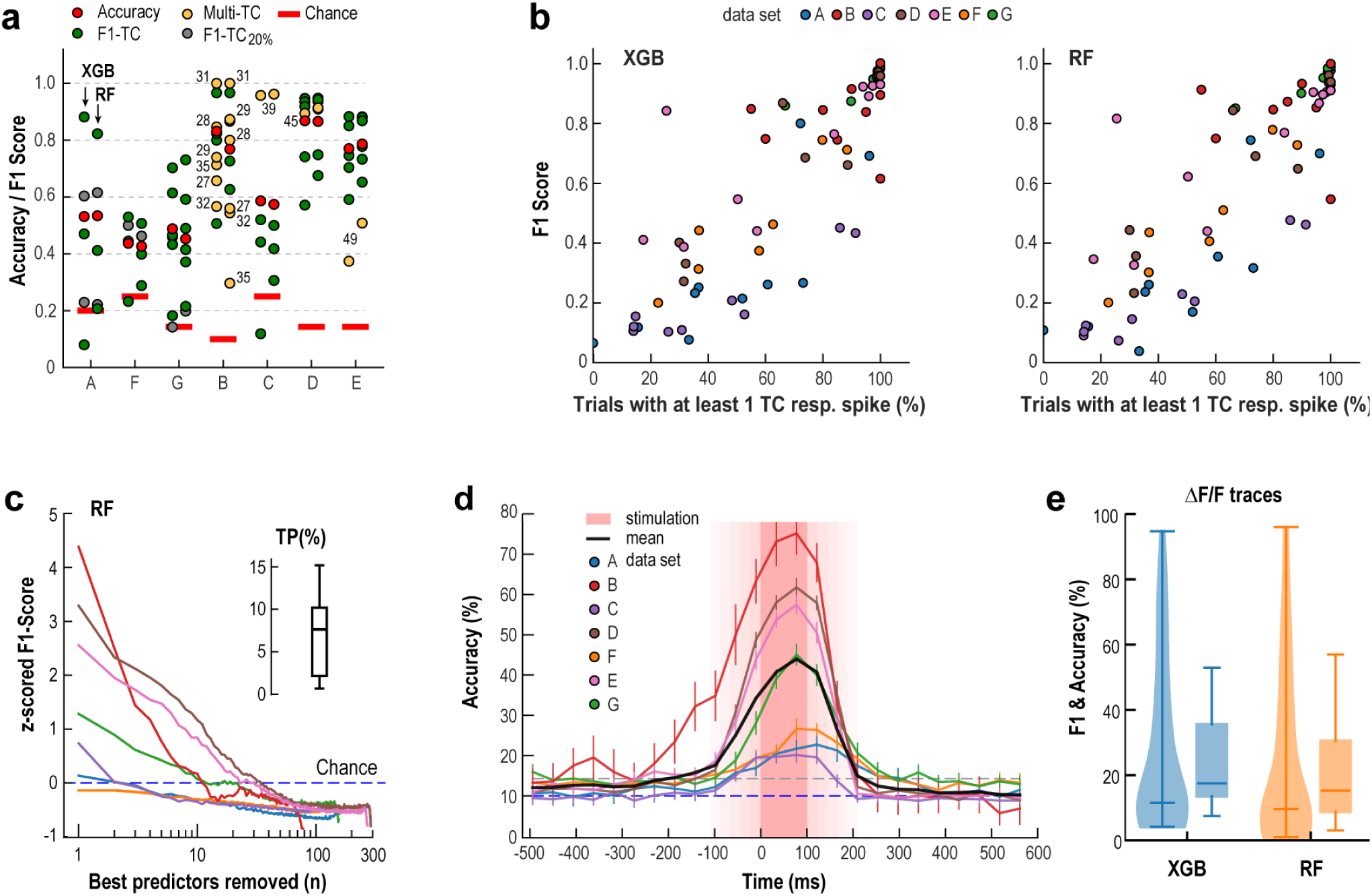
Decoding accuracy, stimulation efficacy and impact of deep interpolation. **a** Overview of F1-scores for TCs (*green*) and TCs that did not reach 20% of spike success (*gray*) upon stimulation. F1-scores are compared to chance level (*red bars*) and overall accuracy (*red circles*). XGBoost (XGB; *left*) and Random Forest (RF; *right*) results are very similar across all datasets. Potential multi-cell targets (cp. Supplementary Fig. 5; TCs 27-29, 31-32, 35, 39, 45 and 49) are labeled and marked in *yellow*, and overall do not show above-average decoding success. **b** For each data set, stimulation success (at least 1 spike) varied among TC. Target stimulation success positively correlated with decoding power, i.e. F1-score, of a TC in XGB (*left*) and RF (*right*). **c** Z-scored F1-scores calculated with the Random Forest method for each TC and then averaged, for each dataset (same color code as in Fig. 4), as function of number of best predictors dropped (cp. XGB in Fig. 4c, *right*). *Inset*: Average number of best predictors dropped before F1-scores reach chance levels. **d** Deep Interpolation slightly advances stimulus information and significantly increases accuracy thereby slightly prolonging transient response time. Analysis done over 132-ms window (cp. Fig. 4f, 22-ms window). The enhanced accuracy achieved with Deep Interpolation demonstrates the method’s noise-reducing benefits, which improve the model’s ability to capture relevant neural signals. However, it also shows that it undermines our ability to study temporal progression of activations, as its window of smoothing ranges across 30 frames. **e** Low (but still above chance) F1 and accuracy when XGB and RF are trained on the raw ΔF/F traces.

**Supplementary Fig. 11.**
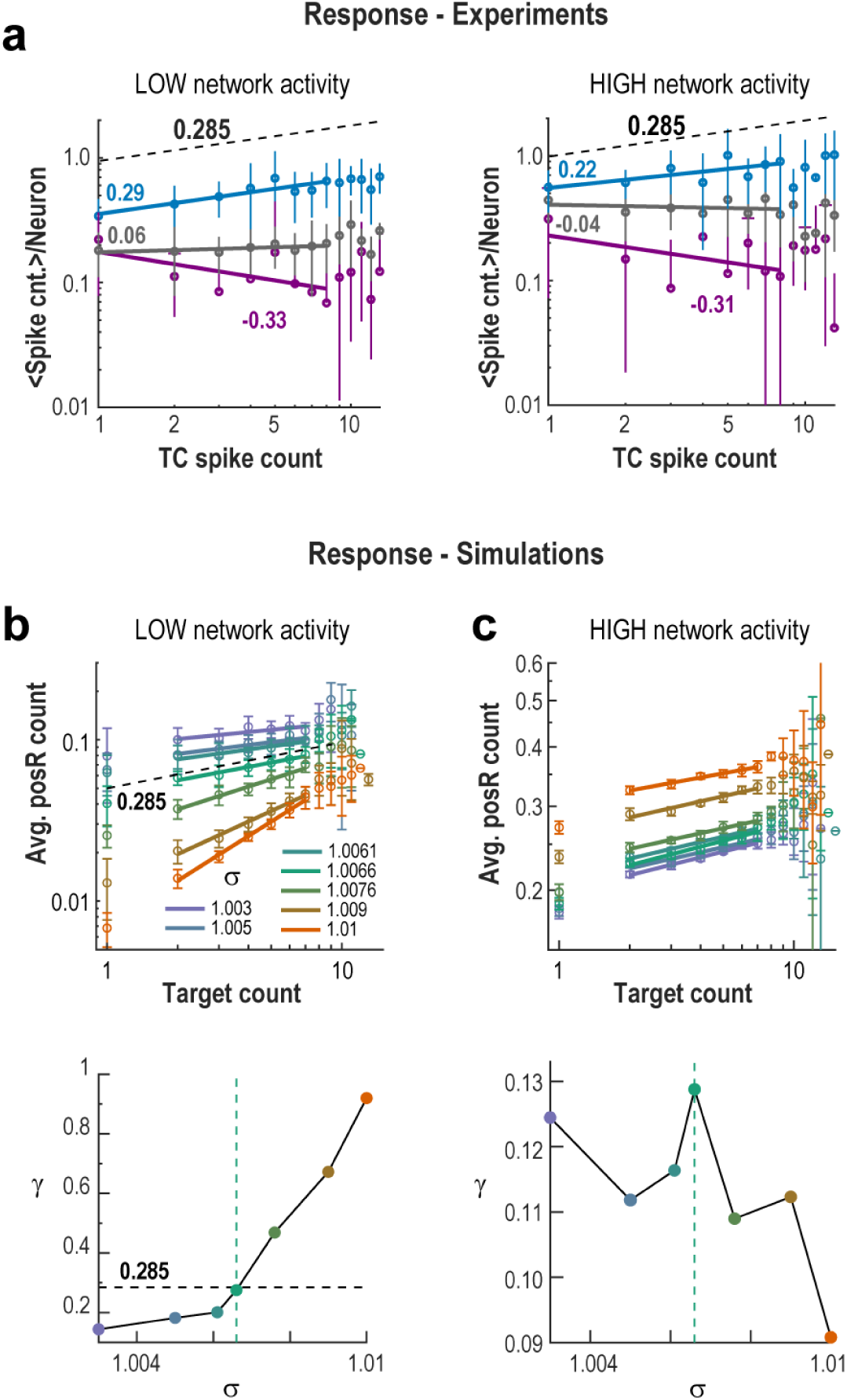
Spike count in subpopulations scale similar with spike count of target cells during low and high avalanche activity states. **a** Mean spike count on PosR (*blue*), NonR (*grey*) and NegR (*purple*) as function of spike count of TC, in log-log, during holographic stimulation. Power law fits are indicated by solid lines, with obtained exponents shown. Separating trials with preceding low (*left*) or high (*right*) baseline network activity. Experimental data. **b** Mean response of PosR as function of TC spike count during trials with low base activity for the simulations. *Bottom*: Response scaling exponents as function of branching parameter σ. **c** Same as in b, but for trials with high base activity prior to stimulation.

**Supplementary Fig. 12.**
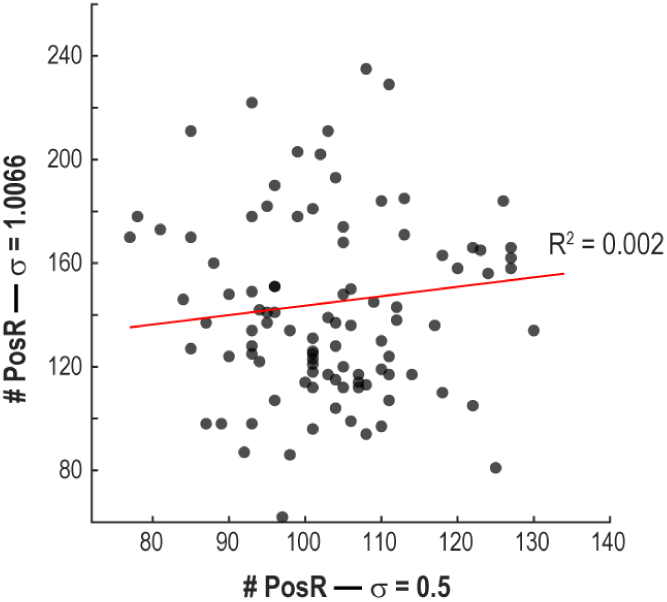
No correlation between number of PosR for TCs between sub- and critical networks with identical connection structure. Number of PosR for each TC (*circles*) at the critical network (σ = 1.0066) as function of the same measure from a subcritical network (σ = 0.5). From the *t*-statistics of the two-sided hypothesis test we obtained p = 0.27 for the slope of the linear regression (*red line*), with R^2^ = 0.002.

**Supplementary Table 1.**

| Experimental response scaling exponents ($\pm 95\%$ C.I.) | | | | |
| --- | --- | --- | --- | --- |
| Condition | Figure | PosR | NonR | NegR |
| Base (neuron) | S6a *left* | $0.46 \pm 0.07$ | $0.52 \pm 0.27$ | $0.44 \pm 0.02$ |
| Base (network) | 2d, S6a *right* | $0.55 \pm 0.22$ | $0.41 \pm 0.27$ | $0.76 \pm 0.24$ |
| Resp (neuron) | 2e, S6b *left* | $0.26 \pm 0.09$ | $0.01 \pm 0.10$ | $-0.30 \pm 0.18$ |
| Resp (network) | 2f, S6b *right* | $0.28 \pm 0.10$ | $0.04 \pm 0.09$ | $-0.07 \pm 0.26$ |
| Calcium | S6c | $0.18 \pm 0.04$ | $0.05 \pm 0.04$ | $-0.18 \pm 0.10$ |
| Low | S9 *left* | $0.29 \pm 0.14$ | $0.06 \pm 0.07$ | $-0.33 \pm 0.50$ |
| High | S9 *right* | $0.22 \pm 0.21$ | $-0.04 \pm 0.18$ | $-0.31 \pm 0.66$ |
| No DeepIP | S10a | $0.35 \pm 0.18$ | $0.08 \pm 0.09$ | $-0.78 \pm 1.48$ |

**Supplementary Table 2.**

| Simulation response scaling exponents ($\pm 95\%$ C.I.) | | | |
| --- | --- | --- | --- |
| Branching parameter | Response (Fig. 7b) | Low (Fig. 7c, left) | High (Fig. 7c, right) |
| 0.3 | $0.015 \pm 0.048$ | N/A | N/A |
| 0.5 | $0.045 \pm 0.031$ | N/A | N/A |
| 0.9 | $0.105 \pm 0.038$ | N/A | N/A |
| 0.95 | $0.133 \pm 0.040$ | N/A | N/A |
| 1 | $0.176 \pm 0.112$ | N/A | N/A |
| 1.003 | $0.179 \pm 0.087$ | $0.144 \pm 0.150$ | $0.124 \pm 0.015$ |
| 1.005 | $0.166 \pm 0.044$ | $0.182 \pm 0.127$ | $0.112 \pm 0.043$ |
| 1.0061 | $0.174 \pm 0.032$ | $0.201 \pm 0.115$ | $0.116 \pm 0.019$ |
| 1.0066 | $0.247 \pm 0.086$ | $0.275 \pm 0.080$ | $0.129 \pm 0.039$ |
| 1.0076 | $0.186 \pm 0.082$ | $0.469 \pm 0.111$ | $0.109 \pm 0.028$ |
| 1.009 | $0.198 \pm 0.070$ | $0.672 \pm 0.108$ | $0.112 \pm 0.038$ |
| 1.01 | $0.189 \pm 0.072$ | $0.920 \pm 0.098$ | $0.091 \pm 0.013$ |

